# Tracing the emergence of the novel fluoroquinolone resistance gene *qrtA* in enterococci through environmental reservoirs and pELF-type linear plasmids

**DOI:** 10.64898/2026.03.26.714403

**Authors:** Yusuke Hashimoto, Masato Suzuki, Trung Duc Dao, Ikuro Kasuga, Thi My Hanh Vu, Taichiro Takemura, Haruka Abe, Futoshi Hasebe, Yusuke Tsuda, Takahiro Nomura, Jun Kurushima, Hidetada Hirakawa, Keigo Shibayama, Hoang H. Tran, Michael S. Gilmore, Haruyoshi Tomita

## Abstract

As animal hosts adapted to land, *Enterococcus* species diverged from a common ancestor shared with *Vagococcus* species by acquiring genes essential for survival and transmission in the exposed arid environment. In addition to their intrinsic ruggedness, human-associated enterococci now frequently exhibit multidrug resistance, largely driven by the accumulation of antimicrobial resistance genes (ARGs) on mobile genetic elements (MGEs). However, the source of these resistance elements, their diversity in natural reservoirs, and the risk that they pose remain unclear. To investigate the movement of novel ARGs from the environmental reservoir and the elements that convey them into clinically important enterococcal lineages, we examined environmental vancomycin-resistant enterococci (VRE) circulating in a polluted urban river in Hanoi, Vietnam. Whole-genome sequencing of VRE isolates revealed multidrug-resistance pELF-type linear plasmids harboring a novel fluoroquinolone resistance gene (*qrtA*) encoding a major facilitator superfamily transporter. The *qrtA* gene appears to have originated on *Vagococcus* chromosomes and to have been mobilized by IS*1216E*-associated MGEs. *Vagococcus* genomes show strong associations between ARGs, IS*1216E,* and plasmids related to those in enterococci, and conjugation experiments demonstrated that pELF-type linear plasmids can transfer between *Enterococcus* and *Vagococcus* without major fitness costs, positioning them as plausible vehicles for importing environmental ARGs into enterococci. Reconstruction of pELF-type linear plasmid evolution in the environment, together with database mining of rare *qrtA*-positive clinical isolates from Asia and Europe, suggests that *qrtA*-bearing pELF-type linear plasmids are in an early phase of global dissemination. These findings highlight the need for coordinated international One Health surveillance of ARG and MGE dynamics, including emerging determinants such as *qrtA*, together with proactive interventions to limit their spread.

## Introduction

*Enterococcus* species arose from common ancestors with *Vagococcus* species and before that, *Carnobacteriaceae. Vagococcus* species are primarily associated with aquatic animals as hosts, while enterococci are more closely associated with land animals, and *Carnobacterium* species are mainly psychrophiles^1^. Since diverging from their last common ancestor with vagococci, enterococci have acquired various genes related to cell wall modification, *de novo* purine biosynthesis, and stress responses, which confer the ability to survive and transmit in harsher environments^1^. The more than 80 *Enterococcus* species that have been identified are found not only in the intestinal tracts of all classes of land animals, but likely as a result to be deposited and persist in a variety of environments including in coastal waterways, soils, and on plants^2^. Some *Enterococcus* species are tightly associated with particular hosts or environmental niches, highlighting their adaptability^3^.

*Enterococcus* species, including *Enterococcus faecium* and *Enterococcus faecalis*, occur in clinical settings as opportunistic pathogens that have been on the rise since 1970^4^. They are inherently resistant to many antimicrobials. This is attributable to a combination of intrinsic traits, such as the chromosomally encoded major facilitator superfamily (MFS) efflux pump gene *emeA* in *E. faecalis*, which is known to be partly responsible for fluoroquinolone (FQ) resistance, and acquired antimicrobial resistance (AMR)^4,5^.

Vancomycin resistance emerged near simultaneously in 2 forms in the mid-1980s, VanA– and VanB-types, in clinical isolates of both *E. faecalis* and *E. faecium*, and prominently spread in a highly challenging multidrug-resistant, hospital-associated clade A1 lineage (CC17: clonal complex 17) of *E. faecium* strains to become a global problem^1–3,6–12^. Currently, the World Health Organization has categorized vancomycin-resistant *E. faecium* (VREfm) as a priority 2 pathogen with potent AMR. The immediate sources from which *E. faecalis* and *E. faecium* acquired the operons that mediate *vanA* and *vanB* has yet to be identified.

In 2019, we discovered an <u>e</u>nterococcal linear-form (pELF-type) plasmid highly associated with AMR including vancomycin resistance^13^. The pELF-type linear plasmid has been detected in *E. faecium,* including VREfm isolated from clinical settings worldwide, and is considered a stable vector of ARGs in a broad range of enterococcal hosts^14^. A nosocomial VRE outbreak involving the transmission of pELF-type linear plasmids to multiple enterococcal species has been reported in Japan, highlighting their potential for rapid AMR dissemination^15,16^.

Treatment options for VRE infections are largely limited to oxazolidinones, novel tetracyclines, and lipopeptides; however, pELF-type linear plasmids capable of conferring combined resistance to linezolid and vancomycin, through acquisition of oxazolidinone-resistance genes, have very recently been reported from multiple regions worldwide^17–19^. Beyond AMR, pELF-type linear plasmids termed pELF_USZ, appear to have broader metabolic utility and were detected in *E. faecium* clinical isolates in Switzerland^20^. That plasmid was shown to have acquired and transmitted a carbohydrate utilization gene cluster from *Enterococcus avium* enhancing the intestinal colonization ability of its *E. faecium* host^21^.

The spread of AMR is the “One Health” issue of concern for humans, animals, and the environment^22^. Especially in countries with limited oversight infrastructure, antimicrobial use in medicine, livestock, and aquaculture, as well as inappropriate use can be prevalent and produce significant amounts of waste containing various antimicrobials^23^. In extreme examples, hospital wastewater may be discharged into rivers without adequate treatment, such as in one instance in Vietnam^24,25^. Selective pressures from residual antimicrobials may lead to the widespread proliferation of AMR bacteria^24^. In such environments, interfaces arise where clinically important bacteria intersect with environmental microbial communities, giving rise to environmental resistomes that accumulate diverse ARGs^26^. From such environmental resistomes, the processes by which novel determinants are occasionally recruited into high-risk human-associated lineages, and the dynamics of MGEs at this interface, nevertheless remain poorly understood^27^.

For the management of AMR bacteria and ARG pollution, environmental monitoring is crucial. VRE and *vanA* genes have been employed in wastewater surveillance^28^. Despite the existence of several reports of clinical VRE isolates carrying the pELF-type linear plasmids, important reservoirs of this plasmid in the environment are largely unknown. Here, we undertook a genomic epidemiological study of VRE isolates in an urban polluted river in Hanoi, Vietnam that is heavily impacted by urban and hospital wastewater, with a focus on AMR and the plasmids that carry ARGs. Within this analysis, we found that, in addition to linezolid, vancomycin, erythromycin, and fosfomycin resistances, a novel transferable FQ resistance gene, consistent with a non-clinical, environmental origin, was present on a suite of stepwise-evolved pELF-type linear plasmid variants. These findings indicate that surface waters impacted by anthropogenic wastewater inputs can act as environmental reservoirs and evolutionary hubs that aggregate resistance elements where hospital-associated VRE, broad-host-range pELF-type linear plasmids and environmental bacteria intersect. This provides valuable insights into the acquisition of ARGs from environmental sources, as illustrated by the emergence of the novel FQ resistance transporter gene *qrtA*, its mobilisation onto, and subsequent dissemination via, pELF-type linear plasmids; broad-host-range ARG vectors at the interface between clinical and environmental compartments.

## Results

### Epidemiology of vancomycin-resistant enterococci and environmental prevalence of pELF-type linear plasmids

In 2021, water samples were collected from the Kim Nguu river in Hanoi, Vietnam, to detect VREs. The Kim Nguu river has been reported to be contaminated with ARGs due to the inflow of urban drainage^24^. We obtained 37 isolates of vancomycin-resistant bacteria. Genome sequence analysis using a short-read sequencer (Table S1) identified 33 of the isolates as *E. faecium*, two as *Enterococcus gallinarum*, one as *Enterococcus raffinosus*, and one as *Aerococcus viridans* (Fig. S1). Of the 33 *E. faecium* isolates, 29 had *vanA*-type and eight had *vanB*-type vancomycin resistance gene clusters. Additional ARGs identified in these isolates included aminoglycoside resistance gene clusters *aac(6’)-aph(2’’)* and *aac(6’)-Ii*, erythromycin resistance genes *erm(B)* and *erm(T)*, and tetracycline resistance genes *tet(L)* and *tet(M)*. Further, two co-harbored genes, such as *cfr(D)*, and *poxtA2* which confer resistance against linezolid—an important reserve drug^17^, and *fosB3*, a gene conferring resistance to fosfomycin. The pELF-type linear plasmid was detected in 30 of the 33 *E. faecium* isolates (91%).

Multilocus sequence typing confirmed that the VanA-type VRE isolates belonged to ST117, ST1693, and ST80, whereas the VanB-type VRE isolates belonged to ST78 and ST17. The 37 core genomes were extracted using Roary, and phylogenetic analysis was conducted^29^. *E. faecium*, *E. gallinarum*, and *E. raffinosus* were classified into different clades (Fig. S1). All *E. faecium* isolates were in clade A1 (CC17) but were grouped into different subclades in the core genome analysis (Fig. S2). The detection of clade A1 *E. faecium* suggests that the rivers are contaminated with hospital wastewater. Furthermore, although the VREfm in the Vietnamese environment were genetically diverse, the pELF-type linear plasmid was widespread among them, indicating that a hospital-associated clade A1 *E. faecium* population heavily colonized by pELF-type linear plasmids has spilled over into, and become established within, the river environment.

Based on the ARGs and core genome information, we elucidated the genetic features of the pELF-type linear plasmids through long-read sequencing analyses of several *E. faecium* isolates. We identified five full-length nucleotide sequences in the genome, including those for pELF-type linear plasmids (plasmid size: 85.0–139 kb; Fig. 1A, Fig. S3A, and Table S2). We detected many ARGs in the pELF-type linear plasmid possessed by strain *E. faecium* NUITM-VRE1, which we named pELF_mdr^30^. Using the current database, no ARGs were identified within the other four pELF-type linear plasmids sequenced. Interestingly, pELF_V3 contained a gene cluster for the carbohydrate utilization system harbored by pELF_USZ, at an identical insertion site (Fig. 1A)^20^. Linear plasmids pELF_V3, or pELF_USZ exhibit 91.4% sequence identity over the total plasmid length, and phylogenetic analysis shows that they are distributed in the same cluster, indicating a common origin (Fig. S3B). Strains NUITM-VRE3 and USZ_VRE35_P46 are of different ST types, indicating horizontal transmission of the linear plasmid (Fig. S3B). These experiments showed the existence of not only diverse VREs but also pELF-type linear plasmids with diverse gene content in Vietnamese rivers, and that related pELF-type linear plasmids are now globally distributed.

**Figure 1.**
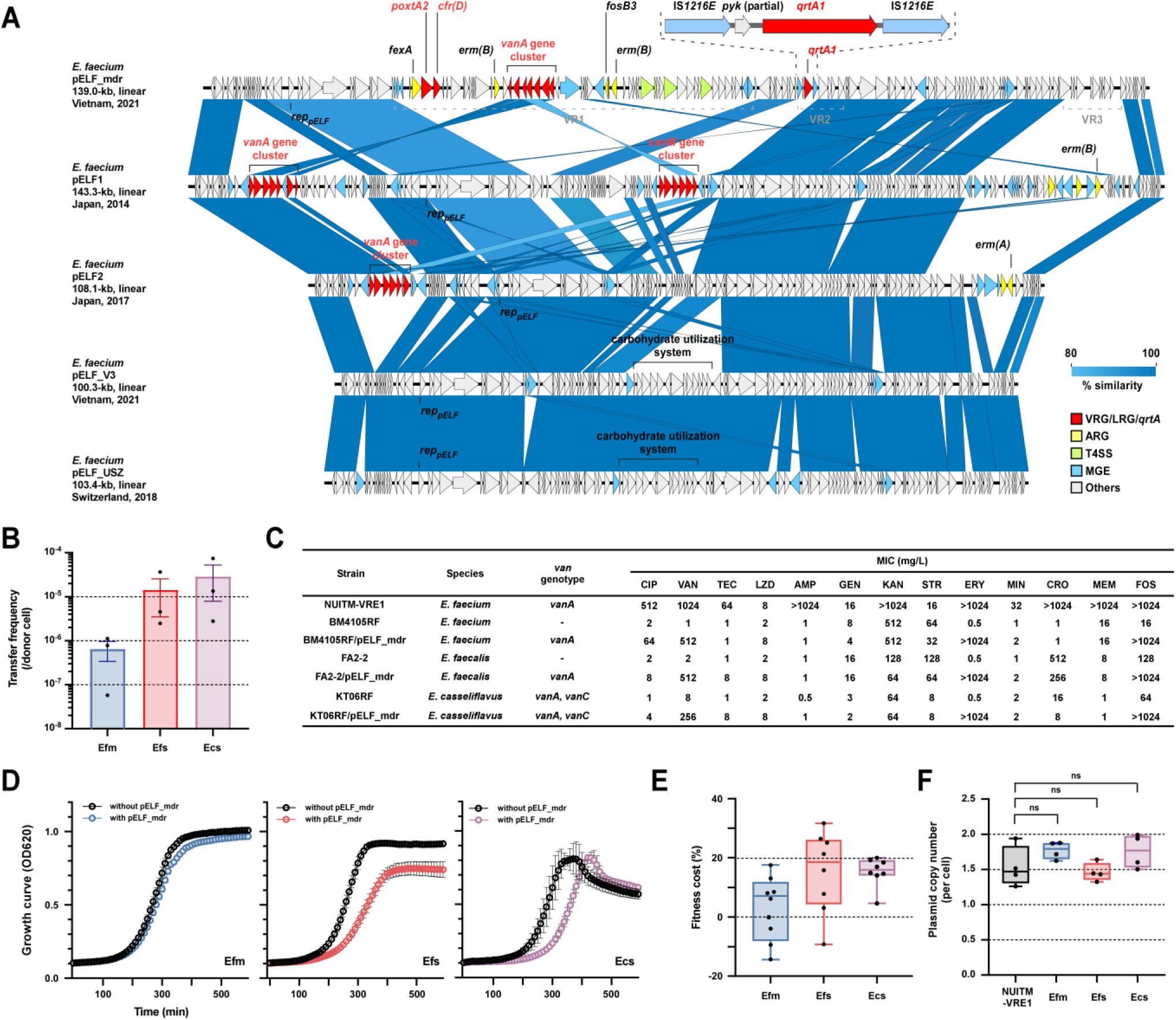
Genetic structures of pELF-type linear plasmids and their effects on host enterococci. (A) Genetic structural comparison of the indicated pELF-type linear plasmids, including pELF_mdr and pELF_V3, in this study. Red arrows show vancomycin resistance gene clusters (VRGs), linezolid resistance genes (LRGs), and *qrtA1* genes; yellow arrows show other antimicrobial resistance genes (ARGs); green arrows show type IV secretion system genes (T4SS)-associated genes; and light blue arrows show mobile genetic elements (MGEs). Variable regions (VR1 to VR3) in pELF_mdr are also shown. (B) Transfer frequencies of pELF_mdr for each recipient cell. Donor strain: *E. faecium* NUITM-VRE1, Recipient strains: *E. faecium* BM4105RF (Efm), *E. faecalis* FA2-2 (Efs), or *E. casseliflavus* KT06RF (Ecs). (C) MICs of the indicated antimicrobials against NUITM-VRE1 and the indicated pELF_mdr-transconjugated enterococci strains. CIP: ciprofloxacin; VAN: vancomycin; TEC: teicoplanin; LZD: linezolid; AMP: ampicillin; GEN: gentamicin; KAN: kanamycin; STR: streptomycin; ERY: erythromycin; MIN: minocycline; CRO: ceftriaxone; MEM: meropenem; FOS; fosfomycin. (D) Growth curves of the indicated pELF_mdr-transconjugated enterococci strains. Each point represents absorbance at OD_620_ measured every 5 min, and the bars represent the standard error of the mean (SEM) (*N* = 8 for each experiment). (E) Fitness costs of the indicated pELF_mdr transconjugant enterococci strains. Whiskers indicate minimum-to-maximum values (*N* = 8). (Welch’s ANOVA, *p* = 0.054). (F) Plasmid copy number of pELF_mdr in NUITM-VRE1 and the indicated pELF_mdr-transconjugated enterococci strains (*N* = 4). ns, not significant (unpaired *t*-test).

Within completely sequenced isolates, NUITM-VRE1, 3, 6, 12, and 30 were found to harbor *vanA* gene clusters on circular T4SS-deficient plasmids with *rep17-type replication* functions, and range in size from 28.5kb to 38.7kb (Fig. S4). In contrast, VanB-type VRE strains NUITM-VRE19 and 32, possessed a Tn*1549*-like element integrated into their chromosomes (Fig. S5).

### Impact of multidrug-resistant plasmid pELF_mdr on *Enterococcus* host

pELF_mdr is a 139-kb linear plasmid with a core structure similar to pELF1, pELF2, and pELF_USZ. It appears to have been formed through acquisition of an *E. faecalis* circular plasmid carrying linezolid resistance genes such as *cfr(D)* and *poxtA2*, together with additional ARGs (Fig. 1A)^30^. Conjugation experiments showed that pELF_mdr transfers to *E. faecium*, *E. faecalis*, and *E. casseliflavus* at frequencies of 10⁻⁵–10⁻⁷ per donor cell, conferring resistance to vancomycin, linezolid, erythromycin, and fosfomycin to transconjugants (Fig. 1B, and C).

In contrast to hospital settings where antimicrobial therapies are common, antimicrobial-selective pressure is thought to be lower in aquatic environments^31^. This raised the question of why plasmids harboring numerous ARGs in such aquatic environments was so abundant. We therefore examined the impact of pELF_mdr, which possesses a wide variety of ARGs, on its host *Enterococcus* with respect to growth parameters and fitness cost^30^. pELF_mdr was found to exert minimal impact on the growth of *E. faecium* and impose almost no fitness cost (Fig. 1D and E). The plasmid copy number remained relatively low, consistent with that found in a previous study of *E. faecium* (Fig. 1F)^14^. In *E. faecalis* and *E. casseliflavus*, the presence of pELF_mdr prolonged the time to log-phase growth and imposed a fitness cost of 10–20% (Fig. 1D and E). Although pELF_mdr harbors regions occurring within *E. faecalis* plasmids, including *rep1*, it imposed a greater fitness cost on *E. faecalis* compared to *E. faecium*. In R2A medium (simulating an aquatic environment), the presence of pELF_mdr did not affect the growth of clade A1 *E. faecium* in either the 25°C or 37°C experimental conditions (Fig. S6). Similar results were observed for strains of both clades A1 and B. In addition to being the predominant plasmid identified among environmental VREs at this sampling site, these results indicate that the pELF_mdr plasmid imposes little fitness cost on the *E. faecium* host and appears to be easily maintained in aquatic environments, supporting a role for pELF_mdr and related pELF-type linear plasmids as stable environmental reservoirs of multidrug resistance.

### Mobilization of a vagococcal MFS-type transporter onto pELF-type linear plasmids

To further explore functions of genes within the *E. faecium*–adapted pELF_mdr plasmid, we explored amino acid sequences and protein structures of gene products on pELF_mdr and identified a novel gene encoding a putative major facilitator superfamily (MFS)-type transporter (Fig. 1A). Given that many MFS-type transporters confer resistance to antimicrobials, we next examined whether pELF_mdr affects drug susceptibility (Fig. 1C). Strikingly, acquisition of pELF_mdr increased resistance to ciprofloxacin in *E. faecium*, *E. faecalis*, and *E. casseliflavus* transconjugants, implicating this MFS-type transporter as a FQ resistance determinant.

All vancomycin-resistant *E. faecium* isolates obtained in this study harbored *efmA*, *efmqnr*, and quinolone resistance–determining region (QRDR) mutations (point mutations in *gyrA* and *par*C), which are known to contribute to FQ resistance (Fig. S1)^32–34^. These FQ resistance factors are chromosomal, and, to the best of our knowledge, no transferable FQ resistance factors have been reported in enterococci. The effect of pELF_mdr was found to be additive when assessed in a clade A1 *E. faecium* background carrying *efmA*, *efmqnr*, and QRDR mutations (Table S3).

We designated this MFS-type transporter gene quinolone-resistant transporter A (*qrtA*) which encodes QrtA—an MFS-type transporter composed of 394 amino acids. To identify remote *qrtA*-like MFS-type transporter homologs, we performed a tBLASTn search, which yielded 4,776 sequences that were subjected to phylogenetic analysis (Fig. 2, and Table S4). The phylogenetic tree resolved six major clusters (Clusters 1–6), each with a largely genus-specific composition. *qrtA* was grouped within Cluster 1 together with sequences from *Enterococcus*, *Vagococcus* and *Lactococcus*, clustering most closely with a *Vagococcus qrtA*-like transporter and with the multidrug efflux pump *emeA* from *E. faecalis*, consistent with *qrtA* belonging to the same lactic-acid–bacterial quinolone-resistance MFS lineage^5^. By contrast, the *Staphylococcus aureus* multidrug efflux pump *norA* grouped in a separate clade together with other staphylococcal homologues (Clusters 3), and well-characterised *Bacillus* multidrug efflux pumps such as *bmr* (*B. subtilis*), *blt* (*B. anthracis*), *ilt* (*B. amyloliquefaciens*) and related *Bacillus* MFS transporters were grouped into distinct clusters (Clusters 2, and 6), indicating that *qrtA* is not a simple derivative of these classical MDR pumps^35–37^. Mapping of gene location showed that most *qrtA*-like homologues are chromosomally encoded, whereas *qrtA* itself is plasmid-borne, highlighting mobilization of this lactic-acid–bacterial transporter from the chromosome onto pELF-type linear plasmids. Thus, pELF-type linear plasmids appear to have captured a transferable FQ resistance determinant from an environmental lactic-acid–bacterial pool, even though, to the best of our knowledge, plasmid-borne FQ resistance has not previously been documented in enterococci.

**Figure 2.**
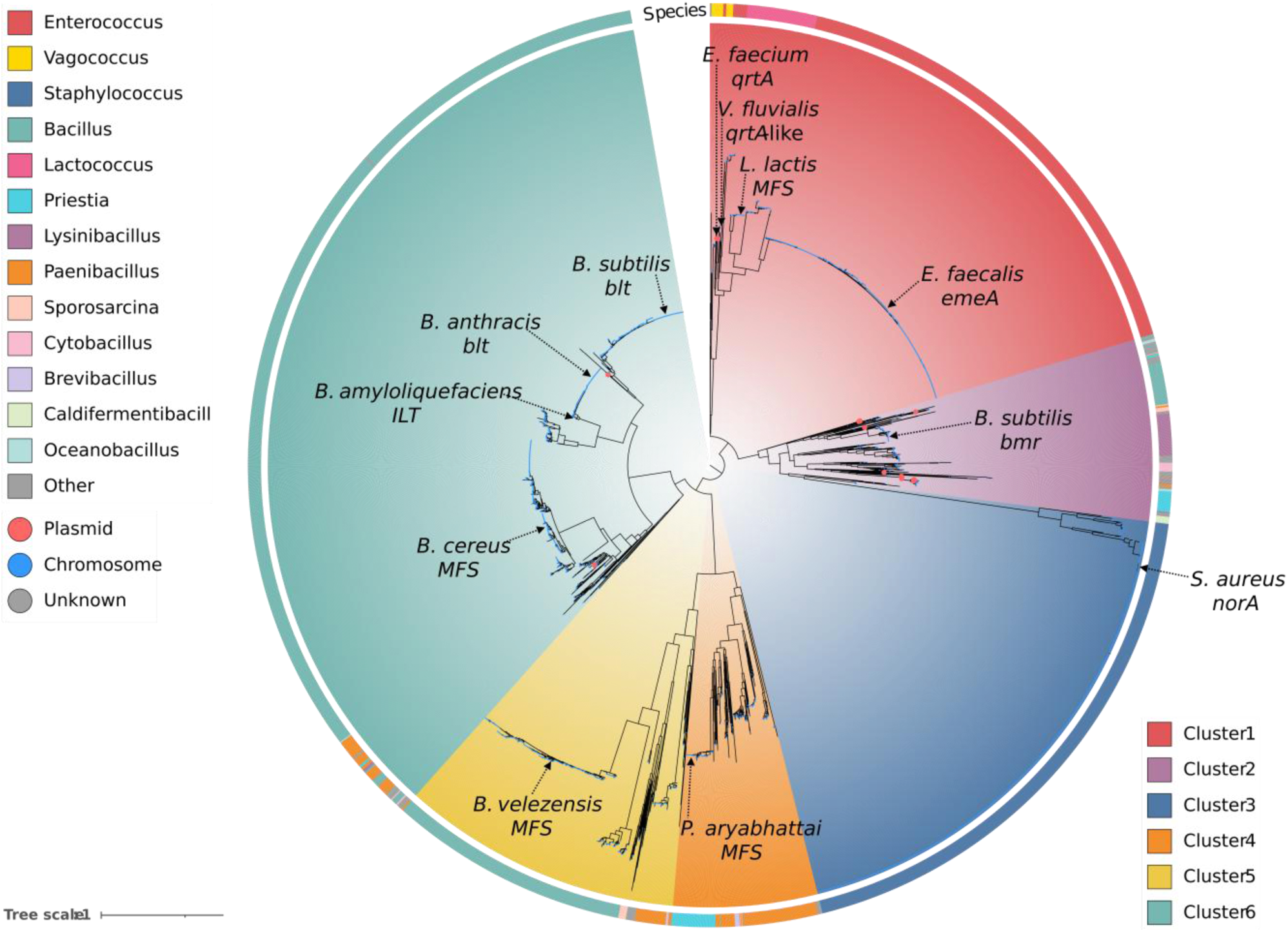
Large-scale phylogenetic tree of MFS-type transporter genes. A phylogenetic tree of *E. faecium qrtA1* and its remote MFS-type transporter homologs was constructed and visualized using iTOL v7. The MFS-type transporter genes were grouped into six major clusters (Clusters 1–6), with representative genes for each cluster indicated by arrows. At the tips of the tree, the genomic location of each gene is shown as plasmid (red circles), chromosome (blue circles), or unknown (black circles). In the outer ring, the bacterial host genus corresponding to each gene is indicated.

### Tracing the origin of transferable fluoroquinolone resistance gene

The MFS-type transporter gene identified on pELF_mdr (*qrtA*) was detected in 25 of the 33 *E. faecium* isolates (Fig. 3A and S1). Because we identified a distinct *qrtA* allele, the pELF_mdr-encoded gene is hereafter referred to as *qrtA1*. One isolate, NUITM-VRE32, carried a distinct allele which we termed *qrtA2a*; *qrtA2a* encodes a product five amino acids shorter and shares 98.7% nucleotide identity with *qrtA1* (Fig. 3A, C, and Fig. S7). Among the isolates for which complete genomes were obtained, pELF_V6 and pELF_V19 also harbor *qrtA1* within the same variable region (VR2) as pELF_mdr, whereas pELF_V12 lacks an MGE containing *qrtA* (Fig. S3A).

**Figure 3.**
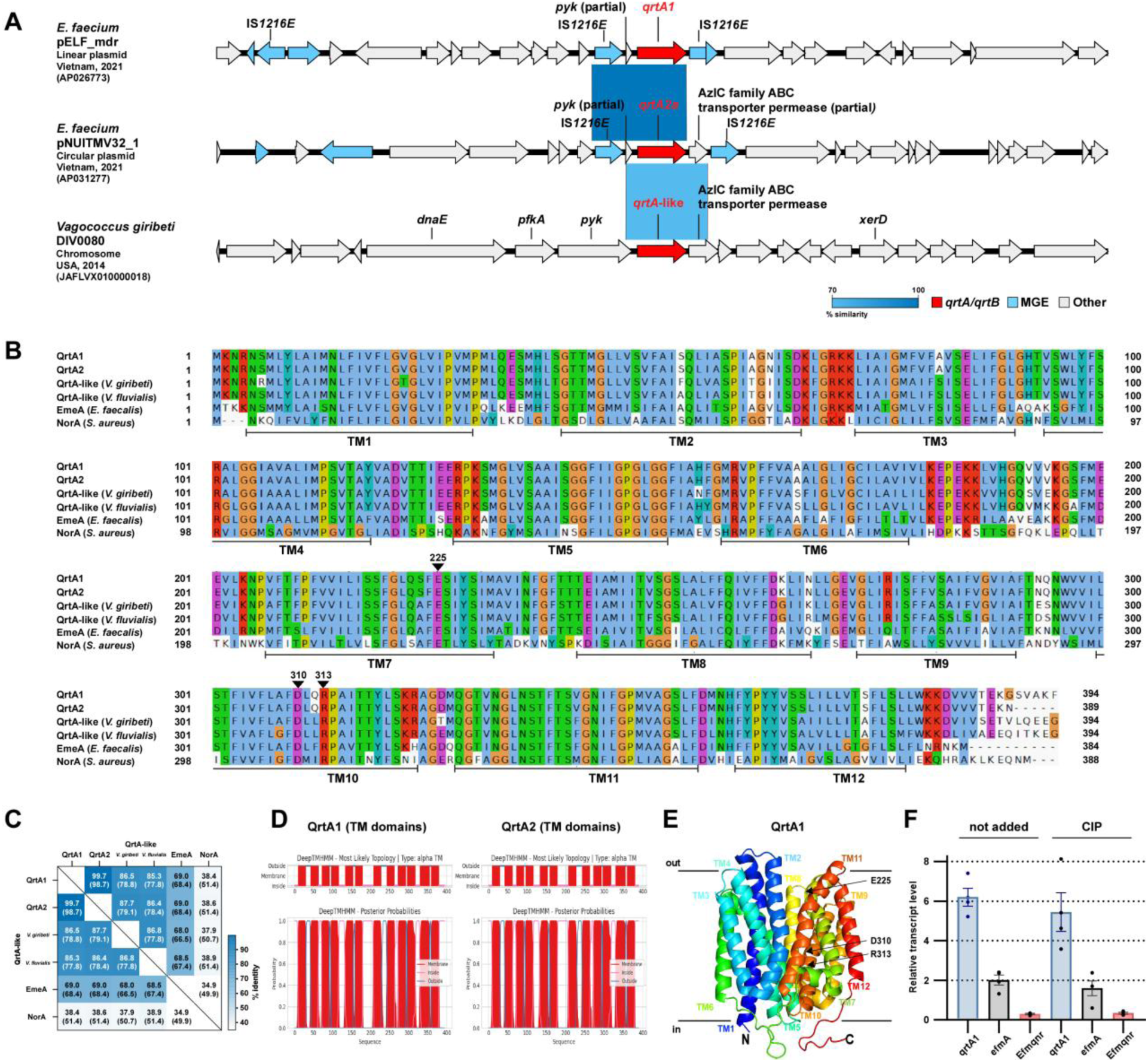
Comparison of MFS-type transporters in Gram-positive bacteria including enterococci and vagococci, and structural features of QrtA. (A) Genetic structural comparison of MFS-type transporter gene-containing regions of pELF_mdr and pNUITMV32_1 in this study, and the chromosome of *V. giribeti* DIV0080. Red arrows show *qrtA*-like genes and light blue arrows show mobile genetic elements (MGEs) including IS*1216E*. (B) Amino acid sequence comparison between QrtA1, QrtA2, QrtA-like gene products from *V. giribeti* DIV0080, and *V. fluvialis* DIV0013, EmeA in *E. faecalis*, and NorA in *S. aureus*. Transmembrane (TM) domains and amino acid residues used in the site-directed mutagenesis are shown. (C) Amino acid sequence identity matrix among the indicated QrtA-like proteins; nucleotide sequence identities are also shown in parentheses. (D) 12 TM domains of QrtA1 and QrtA2. (E) Predicted protein structure model of QrtA1. (F) Transcript levels of the indicated MFS-type transporter gene and other FQ resistance determinants in E*. faecium* NUITM-VRE1. Experiments were conducted without antimicrobials (not added) or with 5 mg/L ciprofloxacin (CIP) (*N* = 4).

Using BLASTp in combination with phylogenetic analysis, we identified related MFS-type transporters from *Vagococcus*, *Lactococcus* and *Enterococcus* spp., including *emeA*, and compared them with *qrtA* (Figs. 3A, B, C, and S8). Upon examining the surrounding genomic structure of the *qrtA1* gene, we identified a partial *pyk* gene upstream of the *qrtA1* gene in pELF_mdr; these genes were flanked by a similarly oriented IS*1216E* element (Fig. S8). A related intact *pyk* gene occurrs on the chromosome of *Vagococcus* spp. Compared to the MFS-type transporter homolog in *Vagococcus giribeti* DIV0080, it exhibits 78.8% nucleotide sequence identity and 86.5% amino acid identity. Comparison of the surrounding structure of the *qrtA*-like gene in the chromosomal genomes of *Vagococcus* DIV0080 and DIV0015 reveals the presence of *pyk* and *qrtA*-like genes in an arrangement similar to the gene cluster on pELF_mdr (Fig. S8).

In addition to the Hanoi river isolates, searches of public genome databases identified *qrtA* variants in four *E. faecium* isolates similar to the NUITM-VRE isolate, and also in one *E. faecalis* strain (Figs. S9 and S10). These five strains were detected in the UK, Belgium, and Malaysia. Among them, *E. faecalis* strain P.En218 was isolated from a human fecal sample, whereas *E. faecium* strains 20160213 and BSAC_ec2084 were isolated from blood samples (Table S5). The nucleotide sequences of *qrtA* genes harbored by these five strains are 100% identical to each other and span 1,370 bases, approximately 15 bp shorter than the 99.6% identical *qrtA1* pELF_mdr gene (Fig. S9). The gene symbol *qrtA*2b was designated for the sequence type identified to occur in the isolates from elsewhere.

Unexpectedly, a pELF-type linear plasmid *rep* (encoding *rep_pELF_*) was identified in the genomes of two *vanB*-type VREs isolated in Belgium (Fig. S10). The genomic assemblies of the five strains harboring the *qrtA2b* gene were characterized at the contig level; hence, the precise loci and adjacent genomic contexts remain undetermined. However, a partial *pyk* gene was identified upstream of *qrtA* in all these strains, and they also shared a gene encoding an AzlC family ABC transporter permease downstream of *qrtA,* the latter being absent in pELF_mdr (Fig. S9). By comparing the upstream and downstream gene structures of *qrtA1* and *qrtA2a*, we infer that IS*1216E* inserted downstream of *qrtA2a* to generate *qrtA1* (Fig. 3A and S8), which further affected the QrtA C-terminus (Fig. 3B). Taken together, these genomic, plasmid-structural and phylogenetic data suggest that we are observing an early stage in the spread of *qrtA* from vagococcal chromosomes into pELF-type linear plasmids circulating in enterococcal popluations including VRE.

Structurally, BLAST comparison indicates that QrtA shares 69.0% amino acid identity with EmeA (Fig. 3C)^38^ with 12 transmembrane domains predicted by DeepTMHMM (Fig. 3D)^39,40^. Comparative structural analysis with AlphaFold2 against NorA and EmeA produced low root mean square deviation scores of 1.9 and 2.0, respectively, suggesting structural similarity (Figs. 3E and S11). Unlike the sequence and structural similarities, the surrounding genomic structures of these MFS-type transporters were quite different (Fig. S8).

We compared transcription levels of the *qrtA* gene in NUITM-VRE1, along with *efmA* and *efmqnr*, which contribute to ciprofloxacin resistance (Fig. 3F), and found that *qrtA* showed the highest transcript levels, which did not change significantly following the addition of ciprofloxacin (Fig. 3F). This indicates that under the conditions tested, *qrtA* is highly expressed constitutively rather than being strongly induced by FQs.

### Functional analysis of *qrtA* gene and impact of transporter inhibitors

The transfer of pELF_mdr led to elevated ciprofloxacin MICs in *E. faecium*, *E. casseliflavus*, and *E. faecalis* (Fig. 1C). To prove the role of *qrtA* on FQ susceptibility, we cloned and expressed the gene and examined its contribution in ethidium bromide (EtBr) efflux assays. The *qrtA1* gene was cloned into the nisin-inducible vector pMSP3535 to assess the MICs under inducible conditions^41^. To quantify the contribution of the *qrtA1* gene in the absence of potential functional redundancy of the *E. faecalis* chromosomal multidrug transporter *emeA,* we constructed and used as the host strain FA2-2_*ΔemeA*^5,42^. In the strains expressing the *qrtA1* gene, a four-fold increase in MICs was observed for ciprofloxacin (Fig. 4A and Table S6), while administration of efflux inhibitors blocked this increase.

**Figure 4.**
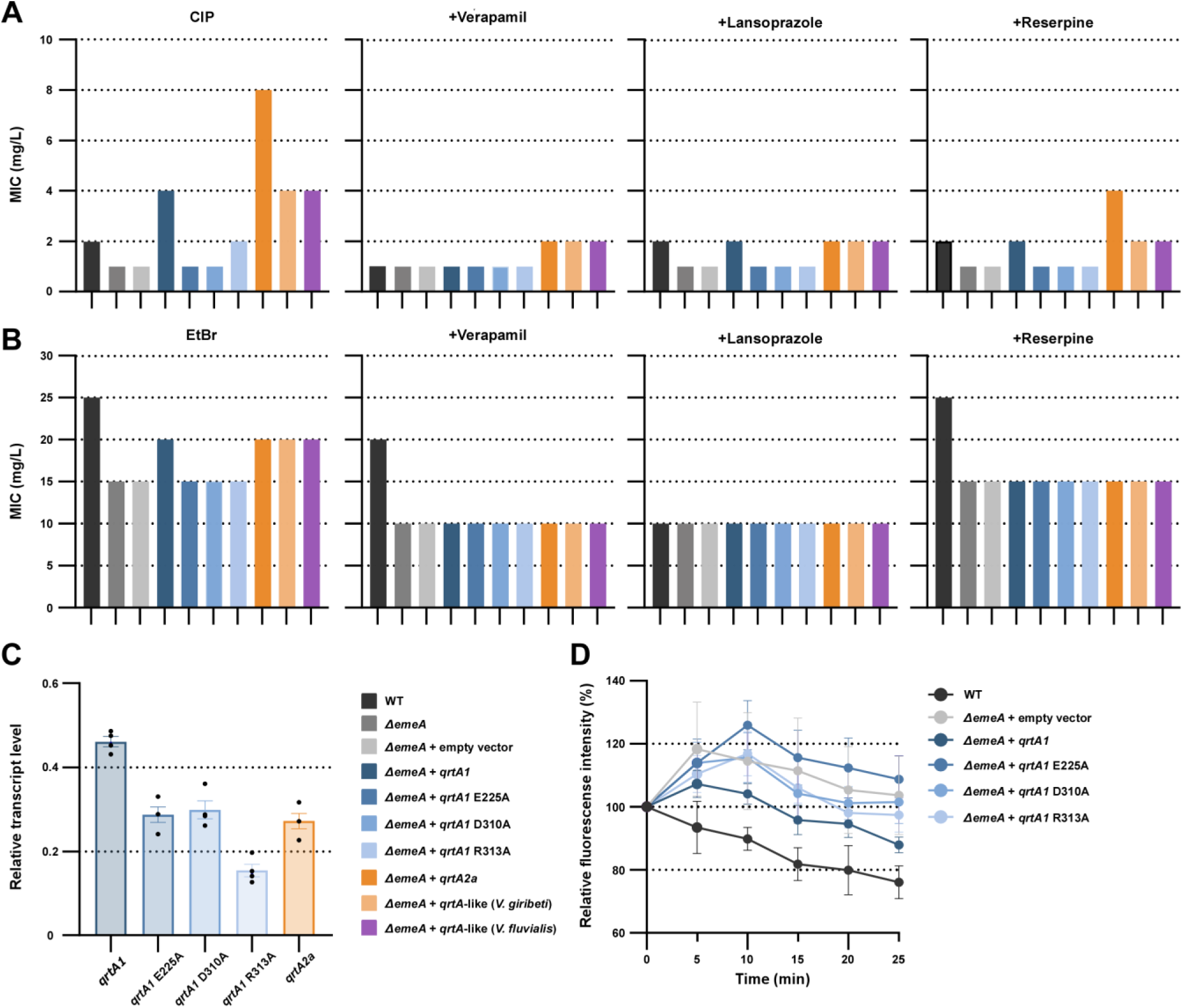
Functional analysis of QrtA-like MFS-type transporters on FQ resistance in enterococci. *qrtA1* and its mutants, and *qrtA*-like genes in enterococci and vagococci were cloned into the expression vector pMSP3535 and introduced into *E. faecalis* FA2-2 (WT) or FA2-2*ΔemeA* (*ΔemeA*) strains. (A, B) MICs of ciprofloxacin (CIP) and ethidium bromide (EtBr) against the indicated *E. faecalis* strains under nisin induction. The experiments were performed in the absence or presence of the indicated efflux pump inhibitors (verapamil, lansoprazole, or reserpine). (C) Transcript levels of the indicated genes in the indicated *E. faecalis* strains (*N* = 4). (D) Efflux capacity for EtBr in the indicated *E. faecalis* strains. The values represent 5-min measurements corrected for the relative fluorescence intensity at the point of initiation and the bars represent the standard error of the mean (SEM) (*N* = 4).

We performed a functional analysis of *qrtA2a* and *qrtA*-like genes in the chromosomes of *Vagococcus* spp. The MICs of the *qrtA*-like genes were equivalent to those of *qrtA1*, and *qrtA2* showed twice the MIC for ciprofloxacin, indicating equivalent functionality to *qrtA1* (Fig. 4A). Experiments with EtBr, which has been reported as an efflux substrate for MFS-type transporters, also showed an increase in MIC values due to the expression of *qrtA1* and its variants (Fig. 4B and Table S6).

Recently, residues critical for NorA transporter function were identified^39^. Four known ionizable residues in the substrate-binding pocket of NorA (Arg98, Glu222, Asp307, and Arg310) form two distinctly charged patches. These residues are also conserved in QrtA1 (Arg101, Glu225, Asp310, and Arg313) (Fig. 3B and E). To determine their impact on ciprofloxacin susceptibility, point mutations were introduced to change the residues E225, D310, and R313 to alanines, amino acids known to affect MIC values in NorA. Using a nisin-inducible plasmid containing cloned genes with E225A, D310A, and R313A mutations, we observed a marked decrease in the MICs for ciprofloxacin and EtBr with the introduction of E225A and D310A mutations in QrtA1, indicating a loss of function (Fig. 4A and B). Introduction of a point mutation to residue R313 resulted in a mild decrease in MIC, unlike in the mutagenesis experiments with the other two residues. These results were consistent with previous results for NorA^39^. The transcript levels of *qrtA1* and its variants in the cloning vector were measured, but we found no correlation with the MIC for ciprofloxacin (Fig. 4C). The low transcript levels of *qrtA2* and *qrtA1* R313A, which had relatively high MICs for ciprofloxacin, suggested that the difference in resistance was dependent on protein function rather than on transcript levels. An EtBr efflux assay was performed to confirm the substrate efflux function of QrtA1^5^. When the *qrtA1* gene was expressed, the relative fluorescence intensity decreased by ∼20% compared to a strain containing an empty vector, indicating active efflux (Fig. 4D). In the *qrtA1* gene with the E225A and D310A mutations, the decrease in fluorescence intensity was abolished such that it could not confer resistance (Fig. 4A).

### pELF_mdr transfer across genera to *Vagococcus* sp

The genus *Vagococcus* is ancestrally related to the genus *Enterococcus*^1,43^. It was first identified in 1989 and is often detected in fish, aquatic animals, and marine and riverine environments^1,44,45^. Human infections with *Vagococcus* spp. are rare; however, there have been some reports of infections caused by *Vagococcus fluvialis* and *Vagococcus lutrae*^46–49^. Little is known of *Vagococcus* plasmid and ARG repertoire.

We performed core genome analysis of *Vagococcus* spp. registered in public databases (Fig. 5A and Table S7). Of the 97 genomes used in the analysis, *qrtA*-like genes were identified in 94.8%, with nucleotide identities ranging from 58.9–78.8% compared to the prototype (Table S7). Vagococcal genomes examined fell into two major clades. The clade containing *V. fluvialis*, which has been reported in association with human infections, carried a *pyk*-*qrtA*-like gene sequence with a relatively high identity to the *qrtA*-like gene (*qrtA*-like gene surrounding structure Type I) (Fig. S12). These findings suggested that an ancestor closely related to this clade was the origin of the *qrtA*.

**Figure 5.**
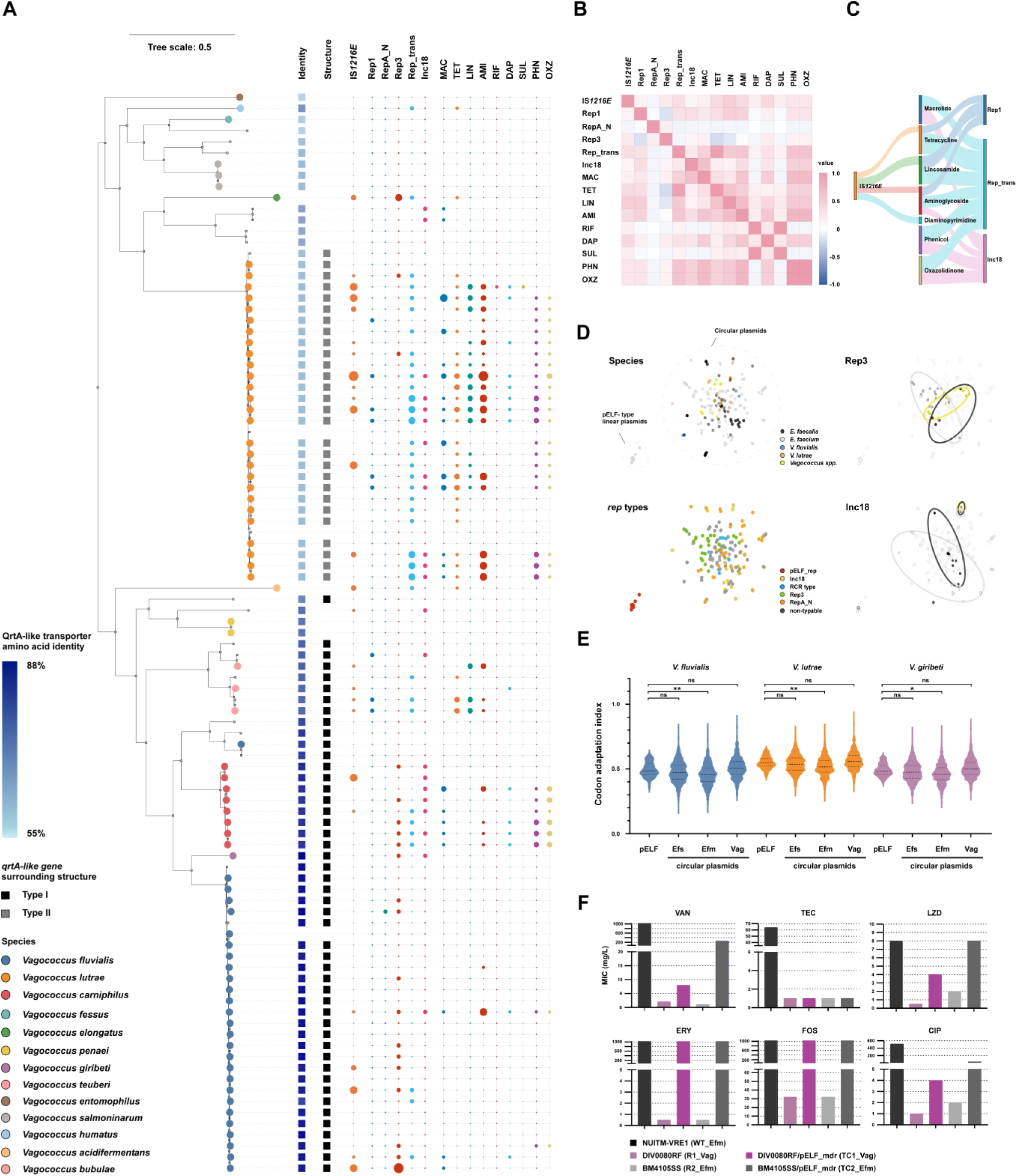
Genomic characteristics of *Vagococcus* spp. and their potential as hosts of pELF-type linear plasmids. (A) Phylogenetic tree of 97 *Vagococcus* spp. strains in the public database, along with genome metadata: bacterial species, *qrtA*-like gene nucleotide identity with *qrtA1*, *qrtA*-like gene-surrounding structure (type 1 or 2), and presence of IS*1216E*, *rep* genes, and ARGs. Dot size reflects copy number (small, single copy; large, multiple copies); absence is not shown. Host species are indicated by colored circle symbols at the tips of the tree. (B) Heat map showing Spearman’s rank correlation coefficients (ρ) for all pairwise combinations, with colours indicating the strength and direction of the association (blue, positive; red, negative; white, no correlation). MAC: macrolide; TET: tetracycline; LIN: lincosamide; AMI: aminoglycoside; RIF: rifamycin; DAP: diaminopyrimidine; SUL: sulfonamide; PHN: phenicol; OXZ: oxazolidinone. (C) Sankey diagram summarizing significant correlations between IS*1216E* (left), ARG classes (middle), and plasmid replication genes (right). The width of each band is proportional to the absolute Spearman correlation coefficient (|ρ|). For each source node, the widths of all outgoing flows add up to 100%. Only associations with |ρ| ≥ 0.3 and q < 0.05 after Benjamini–Hochberg correction are shown in the figure. D) Gene content analysis of pELF-type linear plasmids and conventional circular plasmids of enterococci and vagococci based on the presence or absence of their genes. Bacterial species and plasmid *rep* genes are shown with the indicated color symbols. (E) Codon adaptation index (CAI) of each plasmid population based on the codon usage frequency of the ribosome protein of each species. Blue represents *V. fluvialis*, orange represents *V. lutrae*, and purple represents *V. giribeti*. Efs: *E. faecalis*; Efm: *E. faecium*; Vag: *Vagococcus* spp. ns, not significant; * *p* < 0.05; ** *p* < 0.01 (Dunn’s multiple comparisons test). (F) MICs for the indicated antimicrobials against the indicated donor, recipient, and transconjugants. VAN: vancomycin; TEC: teicoplanin; LZD: linezolid; ERY: erythromycin; FOS: fosfomycin; CIP: ciprofloxacin.

Genomes of strains of the clade including *V. fluvialis* strains harbored few ARGs and plasmid replication functions, whereas strains of the *V. lutrae* clade more frequently isolated from animal sources showed many ARGs and plasmid *rep*s (Fig. 5A). Among these ARGs, *ermB*, *tet(M)*, *aac*(6’)-*aph*(2’), and *ant*(6)-*Ia* were especially frequently detected; ARGs that are common in enterococci including those studied here. High levels of phenicol-resistant *fexA*, *cfrD*, and *optrA* were also observed. These genes appear to be widespread among animal enterococcal isolates and are potentially linked to phenicol use^50^. Many studies have reported the involvement of IS*1216E* in the genomic transition of ARGs in *E. faecium*. When we extended our analysis of mobile genetic elements to vagococci, a search for IS*1216E* in *Vagococcus* identified 26 isolates with full-length copies of this element (Fig. 5A, and Table S7). PlasmidFinder detected 17 plasmid *rep*s within *Vagococcus* genomes (Fig. 5A, and Table S7) including those shared with enterococci. Among the detected *rep* genes, *rep1* belonging to Inc18 and *repUS43* belonging to Rep_trans were particularly abundant and distributed mainly in the *V. lutrae* clade. In the *V. fluvialis* clade, in which relatively rare ARGs were detected, *rep11a* was abundant. No genomes harboring the *rep* gene of the pELF-type linear plasmid were identified. Spearman’s rank correlation analysis revealed strong positive associations between IS*1216E*, multiple ARG classes, and specific *rep* genes in *Vagococcus* genomes (Fig. 5B, C). Macrolide, tetracycline and lincosamide resistance genes were strongly associated with Rep1/Rep_trans, whereas phenicol and oxazolidinone resistance genes were predominantly linked to Inc18-type replicons. Aminoglycoside and diaminopyrimidine resistance genes also correlated with Rep_trans, indicating that these replicons preferentially carry multidrug resistance. Collectively, these data indicate a tight linkage between IS*1216E*, ARGs, and specific plasmid backbones within vagococcal genomes.

To assess similarity between plasmids, plasmidome analysis was performed using 38 full-length plasmid sequences of *Vagococcus* spp. along with plasmids of *Enterococcus* spp. (Fig. 5D). A comparison of the genetic content of the circular plasmids of *E. faecalis*, *E. faecium*, and *Vagococcus* spp. revealed similarities in plasmid content and that the pELF-type linear plasmid formed a single cluster independent of the circular plasmid (Fig. 5D). Examination of the Inc18 and Rep3-family *rep*s and plasmids in *Vagococcus* revealed similarities in plasmid content. In particular, plasmids with Inc18-type replication functions shared 98.3% nucleotide sequence identity within *rep* and 60.2% of the total plasmid length was identical between pBN31-cfrD of *V. lutrae* and pEFS36-2 of *E. faecalis* (Fig. S13A)*. Enterococcus* and *Vagococcus* strains isolated from pig and beef harbored similar plasmids in the Rep3 and Rep2 families (Fig. S13A and B). These findings indicate that enterococci and vagococci share plasmids in nature.

Differences in codon usage frequency between chromosomes and MGEs affect translation efficiency and are related to the fitness cost of MGEs^51^. In all three *Vagococcus* species examined (*V. fluvialis*, *V. lutrae* and *V. giribeti*), Dunn’s multiple-comparison test showed that the Codon Adaptation Index (CAI) distributions of pELF-encoded genes were indistinguishable from those of *Vagococcus* plasmids and *E. faecalis* plasmids (adjusted *p* > 0.99 in all species), but were consistently higher than those of *E. faecium* plasmids (adjusted *p* < 0.05). These findings indicate that the pELF-type linear plasmid is codon-adapted to a level comparable to resident plasmids and better matched to a vagococcal than typical *E. faecium* plasmids, suggesting a relatively low translational cost for maintaining pELF-type linear plasmids in this genus (Fig. 5E).

Since the origin of the *qrtA* gene appears to be or proximal to *Vagococcus*, we investigated the possibility of inter-genus horizontal gene transfer between *Vagococcus* and *Enterococcus* spp. We prepared rifampicin– and fusidic acid-resistant *V. giribeti* DIV0080^43^, since this strain was closely related to *V. fluvialis* and possessed a *qrtA*-like gene with the highest nucleotide identity to *qrtA1* (Fig. 3A). Using this strain as the recipient, we performed a conjugation experiment with NUITM-VRE1, selecting for acquisition of the *vanA* gene, and confirmed the transfer of pELF_mdr (TC1_Vag) (Fig. S14). We also attempted a back cross of pELF_mdr from *V. giribeti* to *E. faecium,* and obtained a secondary transconjugant in the latter host (TC2_Efm). These results indicated that the pELF-type linear plasmid can be transferred between the related genera *Enterococcus* and *Vagococcus.* No differences were observed in the log-phase growth parameter of the recipient and transconjugant strains (Fig. S15). Transconjugant strains showed elevated MICs for vancomycin, linezolid, erythromycin, fosfomycin, and ciprofloxacin, indicating that all of the ARGs are functional in *Vagococcus* (Fig. 5F). The bidirectional transfer and functional expression support the view that vagococci can act both as sources and recipients of pELF-type linear plasmids, thereby linking environmental lactic-acid–bacterial reservoirs with clinically high-risk enterococcal lineages.

## Discussion

Wastewater is an important reservoir of antimicrobials, antimicrobial-resistant bacteria, and environmental microbes^31,52^. Medical, agricultural, and treatment-plant wastewater are hotspots for the acquisition of novel ARGs in clinically important pathogens, as well as for their dissemination^31,53^. Even at concentrations lower than those used clinically, selection pressure from antimicrobials plays an important role in ARG transmission^31,54^. Experiments in activated sludge tanks have shown plasmid– and transposon-mediated resistance transfer in *E. faecalis*, albeit at lower rates than under laboratory conditions^55^. Some MGEs exhibited lower transfer frequencies than the pELF-type linear plasmid *in vitro*, indicating considerable potential for pELF-type linear plasmids to disseminate in the environment. Our genomic analyses now show that this potential has already been realized; a small urban river in Hanoi has become a reservoir where clade A1 VREfm and pELF-type linear plasmids are established.

The pivotal discovery in this study is the identification of the mobilizable *qrtA* gene on the pELF-type linear plasmid, and demonstration of its contribution to FQ resistance in enterococci. *qrtA* encodes a MFS-type transporter that is related to known FQ resistance determinants such as *emeA* in *E. faecalis*, but, in contrast to those chromosomal systems, is carried on a transferable plasmid that has not previously been reported in enterococci. Acquisition of *qrtA* increased ciprofloxacin MICs in *E. faecium*, *E. faecalis*, *E. casseliflavus* and *Vagococcus* sp., and had an additive effect on pre-existing resistance mechanisms mediated by QRDR mutations, *efmA* and *efmqnr*, significantly elevating FQ MICs (Fig. 5F, and Table S3).

FQs are used to treat a wide range of bacterial infections, such as urinary tract and respiratory tract infections. Although FQs are not routinely used to treat enterococcal infections, the 2019 global analysis of the burden of AMR estimated that *E. faecium* resistant to at least one antibiotic caused 51,500 deaths in 2019, including 37,200 attributable to FQ resistance and 14,300 to vancomycin resistance^32,56,57^. This indicate that the global AMR burden attributable to enterococci is driven not only by vancomycin resistance but also by FQ resistance. Furthermore, FQs use has been associated with VRE prevalence, and with increased healthcare-associated infections due to VRE^58,59^. In Vietnam, FQs are widely used in clinical medicine and aquaculture and are frequently detected in wastewater^25,60^. Because they are relatively stable in water and sediments, they are thought to be important drivers of selection for FQ resistance genes^61–63^. Our data support a scenario in which exogenous acquisition of *qrtA* provides an advantage under low environmental FQ selection and further boosts resistance in intestinal populations exposed to higher FQ levels (Fig. 1C and Table S3)^64,65^. In the case of ultra-multidrug-resistant plasmids, such as pELF_mdr, such co-selection will accelerate the spread of multiple resistance traits^66^.

Based on the presence of remnants of flanking genes, gene sequence identity and surrounding genomic structures, our data are most consistent with *qrtA* having originated in *Vagococcus* spp. (Fig. 2, 3A, and 3C). IS*1216E* is consistently present at the end of the *qrtA* MGE supporting its role in mobilizing *qrtA* onto plasmids. Genomic analysis of 97 *Vagococcus* spp. revealed full-length IS*1216E* in 26 isolates, which suggests that IS*1216E* likely mediated initial chromosomal mobilization events for *qrtA* and potentially other ARGs in vagococci^67^. Analogous to the apparent mobilization of *qrtA*, the presence of a small number of MFS-type transporter genes located on plasmids in *Bacillus* strains belonging to clusters 2 and 6 suggests that this mobilization step, although seemingly rare, can indeed occur in different contexts (Fig. 2).

*Vagococcus* spp. share a common ancestor with *Enterococcus* but are not typically commensal bacteria of humans^1^. Despite these ecological differences, genomic analysis of vagococci showed strong positive associations between ARGs, IS*1216E* and plasmids closely related to those in enterococci, indicating a partially shared the mobilome and resistome between species of both genera (Fig. 5A-5D). This supports a model in which vagococci act as reservoirs in some environments in which ARGs can be assembled on IS*1216E*-associated plasmids that are compatible with enterococcal hosts.

Although a pELF-type linear plasmid *rep* gene was not detected in the limited available *Vagococcus* genomic data, our conjugation experiments demonstrated that pELF_mdr can transfer from *E. faecium* to *Vagococcus.* A transconjugant showed no alteration in the growth of *Vagococcus* sp. and was also capable of back transfer, consistent with our CAI analysis indicating that pELF-encoded genes are well codon-adapted to vagococci (Figs. 5E, S14 and S15). pELF-type linear plasmids harbour multiple defense and anti-defense systems potentially contributing to their stability in diverse hosts^30^. Previous work suggested that the broad host range of the pELF-type linear plasmid promoted its transfer to *E. avium*, where it acquired a carbohydrate utilization system gene cluster^20^. Together with our data, this supports a stepwise model in which pELF-type linear plasmids first captured *qrtA* from vagococcal chromosomes and then disseminated this determinant within enterococci. Consistent with this, *qrtA* is now widely disseminated across Vietnam from north to south, being detected in multiple environmental and wastewater samples from climatically distinct regions, including Hanoi and Ho Chi Minh City in 2025, as well as in clinical enterococcal isolates, at least in Hanoi in the same year (data not shown). Given this extensive spread, the identity of the original plasmid that captured *qrtA* cannot be resolved from currently available data. However, complete plasmid sequences begin to clarify the evolutionary steps by which pELF_mdr acquired ARG. pELF_V12, which lacks ARGs is closely related to *qrtA1*-positive plasmids pELF_V6 and pELF_V19 (95.6% nucleotide identity to pELF_V12), consistent with acquisition of an MGE comprising a partial *pyk*–*qrtA1* region and IS*1216E* into pELF_V12 (Fig. S3).

Within the Hanoi river related pELF variants occur in genetically diverse clade A1 VREfm hosts, implying frequent plasmid transfer and establishment of a pELF-type linear plasmid pool that serves as an environmental reservoir of ARGs (Figs. S1–S3). In addition, pELF_V3 was identified with high sequence similarity and core-gene relatedness to the intestinal colonization plasmid pELF_USZ. Thus, multiple stepwise-related pELF-type linear plasmid variants with varying gene content co-occur in this Hanoi river system. At the same time, the close relationship between pELF_V3 and the clinical plasmid pELF_USZ, together with reports of related pELF-type linear plasmids from distant regions, indicates that locally evolved pELF-type linear plasmid variants can join and circulate within a global enterococcal plasmid network^14^.

Variants of the *qrtA1* gene have been found in clinical entecococcal isolates from Asia and Europe. Among these, *rep_pELF_* was identified in the genome of *E. faecium* detected in Belgium in 2016 and 2019 (Figs. S8 and S9). Although the precise location of the *qrtA* gene is unknown because of the contig level of its genome assembly, the presence of *rep_pELF_* suggests that pELF-type linear plasmids in those isolates may already carry *qrtA*. In contrast to the high *qrtA* carriage rates in VRE from the Hanoi river, *qrtA* is extremely rare in publicly available *E. faecium* genomes, consistent with relatively recent dissemination. Our data therefore likely capture an early stage in the emergence of a new plasmid-borne FQ resistance determinant in enterococci and related genera.

Despite the advances made, our study has some limitations. The cross-sectional nature of our sampling limited our ability to assess temporal trends in the spread of pELF-type linear plasmids and the associated resistance genes. There are few studies on VRE reservoirs in the environment. Our study adds to that but admittedly focused on a single geographical region, and further research is required to determine the prevalence and dynamics of these plasmids and *qrtA* genes in other parts of the world as well as in environments less impacted by human activity. Nevertheless, our subsequent surveillance is providing evidence that the distribution of *qrtA* may be broader than even that captured in the present study. Together, these observations indicate that *qrtA* is disseminating and that its spread is not confined to Vietnam but may extend globally. Furthermore, our reliance on short-read sequencing for some analyses may not have fully captured the complexity of the plasmid structures and their genomic contexts. Finally, factors potentially promoting stability and spread, such as toxin-antitoxin, defense/anti-defense, and bacteriocin systems, are encoded on pELF-type plasmids but yet to be functionally examined^68^. Further study will be essential to fully elucidate the contribution of these unusual plasmids to the spread of AMR.

Overall, our findings illustrate how an environmental bacterial reservoir can donate a novel ARG to a clinically important pathogen lineage via a broad-host-range plasmid, in line with conceptual models that emphasize rare but high-impact evolutionary events in the environmental microbiota. They highlight the critical need for a One Health approach to AMR that recognizes the interconnectedness between clinical settings and environments. The widespread environmental presence of multidrug-resistant enterococci and novel transferable *qrtA* gene reported in this study underscore the importance of integrated surveillance and control strategies that explicitly include environmental reservoirs and their plasmid-mediated links to high-risk clinical lineages.

## Methods

### Bacterial strains and drug susceptibility test

Water samples were collected from the Kim Nguu river in Hanoi, Vietnam, in June 2021. After cultivating in Enterococcosel Broth (Becton Dickinson and Company, Sparks, MD, USA) with 4 mg vancomycin, the bacteria were spread onto Enterococcosel Broth with agar containing 6 mg/L vancomycin and cultivated at 37 °C. Colonies that turned black were considered candidate strains of VRE and stored. Using PCR, the bacterial species were identified, and the types of vancomycin resistance as well as the status of pELF-type linear plasmid possession were confirmed. The PCR primers are listed in Table S8. Except where otherwise specified, enterococci were cultured in Todd Hewitt broth (THB; Difco, Detroit, MI, USA) at 37 °C under constant conditions. The agar dilution method was used to determine the MIC, which was interpreted according to the guidelines of the Clinical and Laboratory Standards Institute (http://clsi.org/).

### Transconjugation of bacterial strains

Transconjugants were acquired through filter mating experiments as previously reported^13^. Transconjugants were selected using vancomycin (5 mg/L), rifampicin (50 mg/L), and fusidic acid (50 mg/L). The donor strains utilized were NUITM_VRE1 (accession number: SAMD00533729), while the recipient strains included FA2-2 (*E. faecalis*; accession number: SAMN00809127), BM4105RF (*E. faecium*; accession number: SAMN09464428), ATCC9790RF (*E. hirae*; accession number: SAMN02604142), KT06RF (*E. casseliflavus*), 572RF (*E. faecium* clade A1), and 1021RF (*E. faecium* clade A1).

### Analysis of growth kinetics and plasmid stability

Growth curves were estimated for bacterial strains cultured overnight in 5 mL THB. The bacterial culture was diluted to an absorbance (OD_595_) of 0.2. After centrifugation, the supernatant was discarded, the cells were washed twice with 1 mL PBS, suspended in PBS, and 3 μL of this solution was mixed with 297 μL THB and incubated at 37 °C. Absorbance (OD_620_) measurements were taken kinetically at intervals of 5 min using a Sunrise plate reader (Tecan, Männedorf, Switzerland) during incubation at 25 °C or 37 °C without any agitation. Similarly, growth kinetic experiments in R2A medium, which mimics an aquatic environment, were performed after washing the bacterial cells twice with PBS. Growth rates and fitness costs were assessed using the Growth Rate Program v5.1 (Bellingham Research Institute, Portland, OR, USA) and a formula derived from Tedim *et al*.^69,70^. Briefly, the growth rate of strains bearing pELF_mdr was normalized against that of strains lacking the pELF-type linear plasmid to establish a relative growth rate. Fitness cost was then calculated as follows: *plasmid fitness cost* (%) = (1 – *relative growth rate*) × 100. In the growth curve and fitness cost analyses, eight biological replicates were used, with each strain analyzed in triplicate. Fitness costs were compared using the Mann–Whitney *U* test.

### Plasmid copy number analysis

The copy numbers of pELF-type linear plasmids were calculated using qPCR as previously described^71^: *plasmid copy number* (per chromosome) = (1 + *Ec*)*Ctc* / (1 + *Ep*)*Ctp* × *Sc*/*Sp*. Here, *Ec* and *Ep* are the efficiencies of chromosome and plasmid qPCR amplification (relative to 1), respectively; *Ctc* and *Ctp* are the threshold cycles of the chromosome and plasmid, respectively; and *Sc* and *Sp* are the amplicon sizes (bp) of the chromosome and plasmid, respectively. To measure the copy number of pELF_mdr, DNA was extracted from the bacterial solution grown to the stationary phase in THB medium supplemented with 5 mg/L vancomycin using an ISOPLANT kit (Nippon Gene, Tokyo, Japan). Considering its distance from *oriC*, the *gyrB* gene was used as the internal control. Plasmid copy numbers were compared using unpaired *t*-test. The primers are listed in Table S8.

### Whole-genome sequencing and genomic analysis

Total DNA was extracted using a Gentra Puregene Yeast/Bact. kit according to the manufacturer’s instructions (QIAGEN, Hilden, Germany). Short reads were generated using the MiniSeq system (Illumina, San Diego, CA, USA) and a High-Output Reagent Kit (300 cycles). Libraries with insert sizes of 500–900 bp were curated using a Nextera XT DNA Library Prep Kit (Illumina). A DNA library was constructed using a SQK-RBK004 Rapid Barcoding Kit according to the manufacturer’s guidelines and library sequencing was performed on a MinION Flow Cell (R9.4.1; Oxford Nanopore Technologies, Oxford, UK). Long reads were assembled *de novo* using the Canu assembler (2.1.1), and subsequent refinement was conducted using short reads in Pilon (1.20.1)^72–74^. For functional annotation, assembly sequences were processed using the Prokka, RAST, and DFAST annotation pipelines^75–77^. For core genome analysis, genome annotation was uniformly conducted using Prokka software. Core genes were identified and aligned using Roary (3.13.0)^78^, from which a phylogenetic tree was constructed using RAxML (8.2.4) under the GTRGAMMA model and visualized using Figtree software (1.4.4)^79^. Phandango, Easyfig, and Genome Matcher (3.0.2) were utilized for data visualization and genome comparisons^80–82^. Multilocus sequence typing, antimicrobial resistance genes, replicon types, toxin–antitoxin systems, virulence genes, and MGEs were identified using various tools, including Staramr (https://github.com/phac-nml/staramr), ResFinder, ABRicate (https://github.com/tseemann/abricate), PlasmidFinder, TASer, VirulenceFinder, ISfinder, and MobileElementFinder^83–88^.

Associations among IS*1216E*, ARG classes (macrolide, tetracycline, lincosamide, aminoglycoside, rifamycin, diaminopyrimidine, sulfomamide, phenicol, oxazolidinone), and plasmid replication genes (Rep1, RepA_N, Rep3, Rep_trans, Inc18) were assessed using Spearman’s rank correlation coefficient (ρ). Variables were encoded as presence/absence, and pairwise complete observations were used. For each pair, two-sided ρ and p values were computed in GraphPad Prism (v8). To control for multiple testing across all pairwise comparisons, p values were adjusted using the Benjamini–Hochberg false discovery rate (FDR) procedure, yielding q values. Statistical significance was defined as q < 0.05, and |ρ| ≥ 0.3 was used as a threshold. The heat map shows all pairwise Spearman ρ values without significance masking on a fixed −1 to +1 diverging scale (blue, positive; red, negative; white, ∼0), with no clustering applied. The Sankey diagram includes links meeting q < 0.05 and |ρ| ≥ 0.3; link weights are set to |ρ| and normalized within each source such that outgoing flows sum to 100%. Plots were generated in Python (Plotly). Plasmid gene network analysis was performed using PANINI from Roary’s output file and visualized using Microreact^89,90^. Codon adaptation index (CAI) values were calculated using the cai program in the EMBOSS package^91^. For each host species, genes encoding ribosomal proteins were used as the reference set to derive codon usage frequencies, and CAI values were computed for all chromosomal and plasmid-borne coding sequences, including those on pELF-type linear plasmids, *E. faecalis*-, *E. faecium*-, and *Vagococcus*-derived plasmids^92^. CAI distributions were compared using the Kruskal–Wallis test followed by Dunn’s multiple-comparison test, with multiplicity-adjusted P values reported (GraphPad Prism (v8)).

### Phylogenetic analysis of *qrtA*–related MFS efflux pump genes

To investigate the phylogenetic relationships of homologs of the MFS efflux pump gene *E. faecium qrtA1* (1,185 bp, accession no. AP026773), TBLASTN searches were conducted against the NCBI Nucleotide collection (nt; accessed 15 June 2025) using amino acids of the encoded protein (394 aa) as the query. Hits with ≥90% query coverage (n = 4,968) were retained. Multiple sequence alignment was generated using MAFFT v7 with the FFT-NS-1 strategy and subsequently trimmed with Gblocks v0.91b with the settings –t=d –b4=5 –b5=h, reducing the alignment length from 1,581 to 924 nucleotide positions (58%) across 18 conserved blocks^93,94^. A maximum-likelihood tree was inferred using FastTree v2.1.11 under the GTR+CAT model with default settings^95^. To remove sequences exhibiting unusually long branch lengths, phylogeny-based clustering was performed using TreeCluster v1.0.3 with the single-linkage method. The threshold value (-t = 1.6) was empirically determined from the pairwise distance between *qrtA1* and the MFS efflux pump gene S. aureus *norA* (1,167 bp, accession no. AB086042), and the largest cluster including *qrtA1* and *norA* (n = 4,776) was retained for downstream analyses^96^. The sequences were realigned using MAFFT v7 and trimmed with Gblocks v0.91b under the same conditions as described above, reducing the alignment length from 1,452 to 959 nucleotide positions (66%) across 8 conserved blocks. A maximum-likelihood tree was inferred using IQ-TREE v2.4.0 under the MFP model with 1,000 ultrafast bootstrap replicates^97^. The final tree consisting of six clusters and associated metadata were visualized using iTOL v7^98^.

### Protein structure prediction

Protein structure prediction was performed using amino acid sequences in AlphaFold 2.3.1 on the Galaxy Europe server with default parameters^99,100^. A structure-based protein homology search using AlphaFold-predicted structural models was performed for the PDB100 database on the FoldSeek server in 3Di/AA mode^101^. AlphaFold-predicted and experimental structures were visualized and compared using PyMOL software 2.5.8 (The PyMOL Molecular Graphics System, Schrödinger, LLC).

### Transcription level measurement

For the transcript level measurement, bacteria were first cultured overnight in Brain Heart Infusion (BHI) medium at 37°C. The overnight cultures were diluted 1,000-fold into fresh BHI medium and grown until the late exponential phase. Cells were treated with RNAprotect Bacterial Reagent (Qiagen) for stabilization, and RNA was extracted from 1 mL of the culture using the RNeasy Minikit (Qiagen), according to the manufacturer’s instructions. For RT-qPCR, cDNA synthesis was carried out using the PrimeScript RT Master Mix (Takara Bio, Shiga, Japan). Real-time PCR was then performed on an ABI 7500 Fast RT-PCR system (Thermo Fisher, Waltham, MA) using Luna Universal qPCR Master Mix (New England BioLabs, Ipswich, MA). The primers are listed in Table S8.

### Preparation of *emeA* gene disruption strains

To analyze the function of the *qrtA* gene, we generated a strain with disrupted *emeA* gene expression (FA2-2*ΔemeA*). To create the FA2-2*ΔemeA* strain, we used the genetic manipulation method with a pCJK47 vector^102^. The DNA fragments to be inserted were constructed by PCR using the respective primers and inserted into the pCJK47 vector using the restriction enzyme *Eco*RI (Roche) and a DNA ligation kit (Takara Bio, Shiga, Japan) as previously described (Table S8)^103^. After transformation into *Escherichia coli* EC1000, the strains were incubated in 5 mL Luria–Bertani (LB) medium containing 300 mg/mL erythromycin at 37 °C with shaking. Recombinant plasmid DNA was extracted using a QIAprep Spin Mini Kit (QIAGEN). After confirming insertion of the desired sequence, electroporation of CK111/pCF10-101 was performed. Overnight cultures of the donor strain (CK111/pCF10-101, pCJK47 derivatives) and recipient strain (FA2-2) were diluted 100-fold in fresh THB broth and incubated separately at 37 °C for 1 h. We mixed 100-µL aliquots of donor and recipient cultures were mixed with 800 μL fresh THB, incubated at 37 °C with shaking at 150 rpm for 20 h, spread on a BHI plate [containing 25 mg/mL rifampicin, 25 mg/mL fusidic acid, 10 mg/mL erythromycin, and 100 mg/mL 5-bromo-4-chloro-3-indolyl-β-D-galactoside (X-gal)], and incubated at 37 °C for 48–72 h. Blue colonies were considered to have successfully integrated the pCJK47 derivative plasmid into the chromosomal target locus and were isolated and purified. These integrant clones were inoculated into THB and incubated at 37 °C for 12 h. After 100-fold dilution, 100 μL of the culture broth was placed onto a MM9YEG plate supplemented with 10 mM 4-chloro-DL-phenylalanine (Sigma-Aldrich, St. Louis, MO, USA) and 40 mg/mL X-gal, and incubated at 37 °C for 12 h. White colonies were isolated and purified as potential candidates carrying mutations within the target genes. Nucleotide sequences were assessed to confirm successful nucleotide substitution.

### Cloning experiments of *qrtA1* and its variants

Cloning experiments were performed to analyze the functions of *qrtA* and its variants using the pMSP3535 vector harboring the nisin-inducible promoter and either the FA2-2 or FA2-2Δ*emeA* strain as host^41^. The DNA fragments were constructed by PCR using the respective primers and inserted into the pMSP3535 vector using the appropriate restriction enzymes and a DNA ligation kit (Takara Bio) (Table S8). After restriction enzyme treatment (New England BioLabs, Ipswich, MA, USA) and ligation, transformation to DH5α was performed and colonies were selected LB supplemented with 125 mg/L erythromycin. Cloning of the target insert was confirmed by Sanger sequencing using the Genetic Analyzer 3130XL (ThermoFisher Scientific, Waltham, MA, USA). Further functional analysis of *qrtA1* was performed through point mutation introduction experiments in E225A, D310A, and R313A (Table S8) using a KOD-Plus-Mutagenesis Kit (TOYOBO, Osaka, Japan). Briefly, pSMP3535::*qrtA1* was used as a template, Inverse PCR was performed with each primer set, followed by treatment of the template DNA with *Dpn* I, and PCR products were circularized by self-ligation (Table S8). Chemical transformation was performed to DH5α, and selection was performed in LB supplemented with 125 mg/L erythromycin. The desired mutation was confirmed using Sanger sequencing.

### Ethidium bromide efflux assay

Bacterial cultures were incubated in THB with 150 mg/L erythromycin at 37 °C overnight. We transferred the culture at a 1:4 ratio to BHI liquid medium containing 100 ng/mL nisin and performed incubation for 2 h at 37 °C. Following incubation, 300 µL of the culture was centrifuged, washed twice with 1 mL 20 mM HEPES buffer (pH 7.0), and resuspended in the same buffer to a volume of 300 µL (OD_620_ = 0.6, ∼2 × 10^8^ CFU/mL). The bacterial suspension was treated with EtBr as efflux substrate (final concentration 2.5 µM) at 37 °C after adding CCCP (final concentration 40 µM) to eliminate membrane potential. After 1 h, the cells were washed thrice with 1 mL HEPES buffer containing 2.5 µM EtBr and resuspended in 3000 µL of the same buffer (∼2 × 10^7^ CFU/mL). The cells were kept on ice until efflux was initiated by adding 8 µL THB to 300 µL of the cell suspension. The efflux pump inhibitors reserpine (6 µg/300 µL), verapamil (30 µg/300 µL), and lansoprazole (30 µg/300 µL) were added prior to the THB. The inhibitor concentrations were selected based on a prior study^5^. Fluorescence measurements were recorded every 30 s for 30 min at excitation and emission wavelengths of 500 and 590 nm, respectively, using an EnSpire spectrophotometer (PerkinElmer, Shelton, CT, USA).

### Pulsed-field gel electrophoresis

Pulsed-field gel electrophoresis was performed as previously described^13^. Briefly, 1% agarose gel blocks containing embedded enterococci were treated with lysozyme solution (10 mg/mL; Roche, Basel, Switzerland) at 37 °C for 6 h, followed by proteinase K solution (60 mAnson U/mL; Merck Millipore, Darmstadt, Germany) at 50 °C for 48 h. To confirm the chromosomal type, the blocks were treated with *Sma*I for 20 h at 25 °C. Blocks without enzyme treatment were used to confirm the linear plasmids. Pulsed-field gel electrophoresis was performed using a CHEF-MAPPER (Bio-Rad Laboratories, Richmond, CA, USA) according to the manufacturer’s instructions (6.0 V/cm, 4 °C, 5.3–66-s pulse, over 19.5 h).

## Acknowledgements

We are grateful to Hideru Obinata and Yoko Sagara (Gunma University Graduate School of Medicine) for helpful discussions and their help regarding ethidium bromide efflux assay and Ryan Whipple (Harvard Medical School) for the valuable technical contributions. This work was supported by grants (JP25fk0108665, JP26fk0108755, and JP25wm0225029 to H. Tomita; JP25fk0108665, JP25fk0108683, JP25fk0108712, JP26fk0108755, JP26fk0108756, JP25gm1610003, JP25wm0225029, JP25wm0225054, JP25wm0325076, and JP25wm0125012 to M. Suzuki; JP25fk0108664, JP25fk0108665, JP25fk0108712, JP26fk0108755, and JP25wm0225029 to I. Kasuga; JP25wm0225029 and JP21wm0125006 to F. Hasebe; JP25wm0225029 to Y. Tsuda; JP25fk0108665, JP25gm1610003, and JP26fk0108755 to K. Shibayama) from the Japan Agency for Medical Research and Development (AMED), Japan, grants (22K16368 to Y. Hashimoto; 22K07067 to H. Tomita; 22KK0058 and 23K06556 to M. Suzuki; 22KK0058 to I. Kasuga; 23KK0151 to K. Shibayama) from the Ministry of Education, Culture, Sports, Science and Technology (MEXT), Japan, grants (21KA1004 and 24KA1005 to H. Tomita) from the Ministry of Health, Labor and Welfare (MHLW), Japan, grants (H. Tomita, M. Suzuki, K. Shibayama) from Consortium for the Exploration of Microbial Functions of Ohsumi Frontier Science Foundation, Japan, a grant (Kambe Memorial Research Award to Y. Hashimoto) from Gunma Foundation for Medicine and Health Science, Japan, a grant (AI083214 to M.S. Gilmore) from National Institutes of Health (NIH), the United States, and a grant (108.02-2017.320) from National Foundation for Science and Technology Development (NAFOSTED), Vietnam.

