## Supplemental figures (Figure S1 to S15) for "Tracing the emergence of the novel fluoroquinolone resistance gene *qrtA* in enterococci through environmental reservoirs and pELF-type linear plasmids"

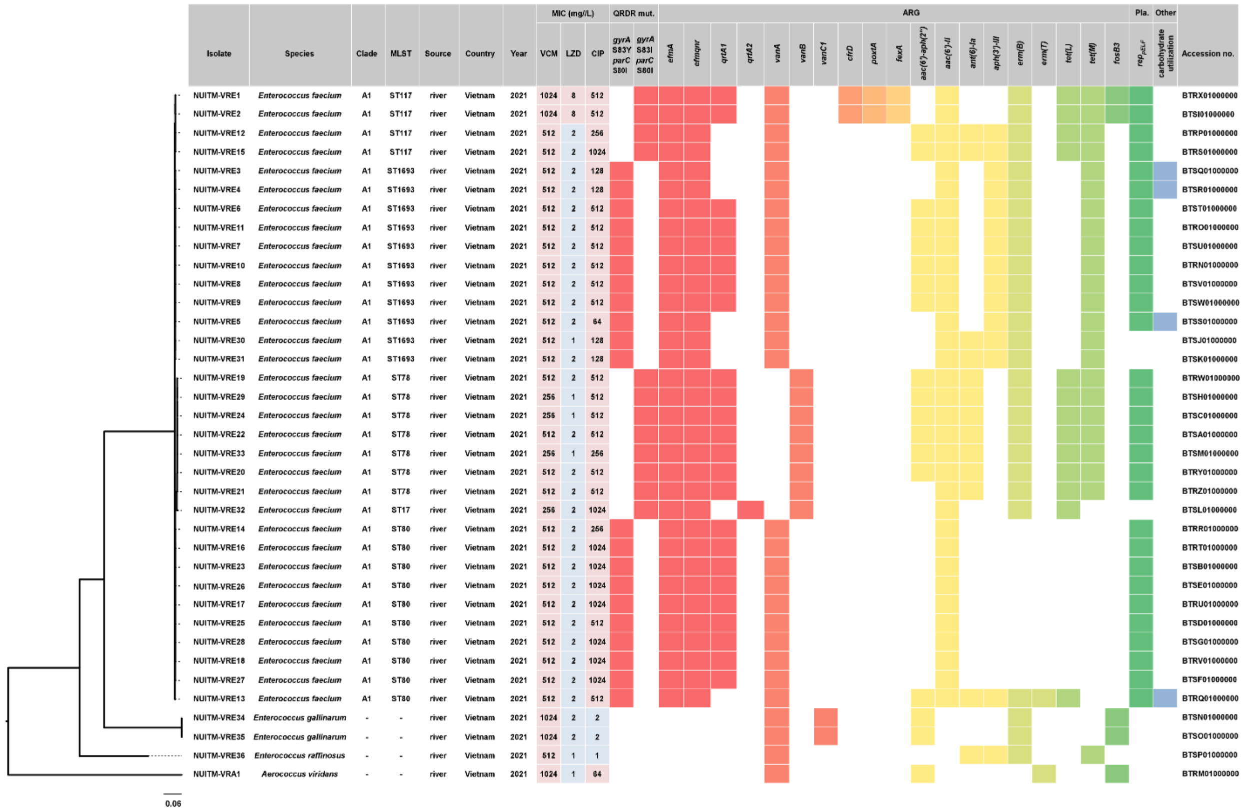


**Figure S1. Phylogenetic analysis and characteristics of vancomycin-resistant bacteria isolated in Vietnam.** Phylogenetic tree of 37 vancomycin-resistant bacterial isolates in this study, bacterial species, source, source of isolation, county of isolation, year of isolation, MICs of the indicated antimicrobials (resistant and susceptible are highlighted in red and blue, respectively), quinolone resistance-determining region (QRDR) mutations, and accession number are shown. The presence of the indicated antimicrobial resistance genes (ARGs), pELF-type linear plasmid *rep* gene ($rep_{\mathrm{pELF}}$), and carbohydrate utilization-related gene cluster are also shown. For *E. faecium* isolates, clade and sequence types (STs) from multilocus sequence typing (MLST) analysis are included. VAN: vancomycin; LZD: linezolid; CIP: ciprofloxacin.

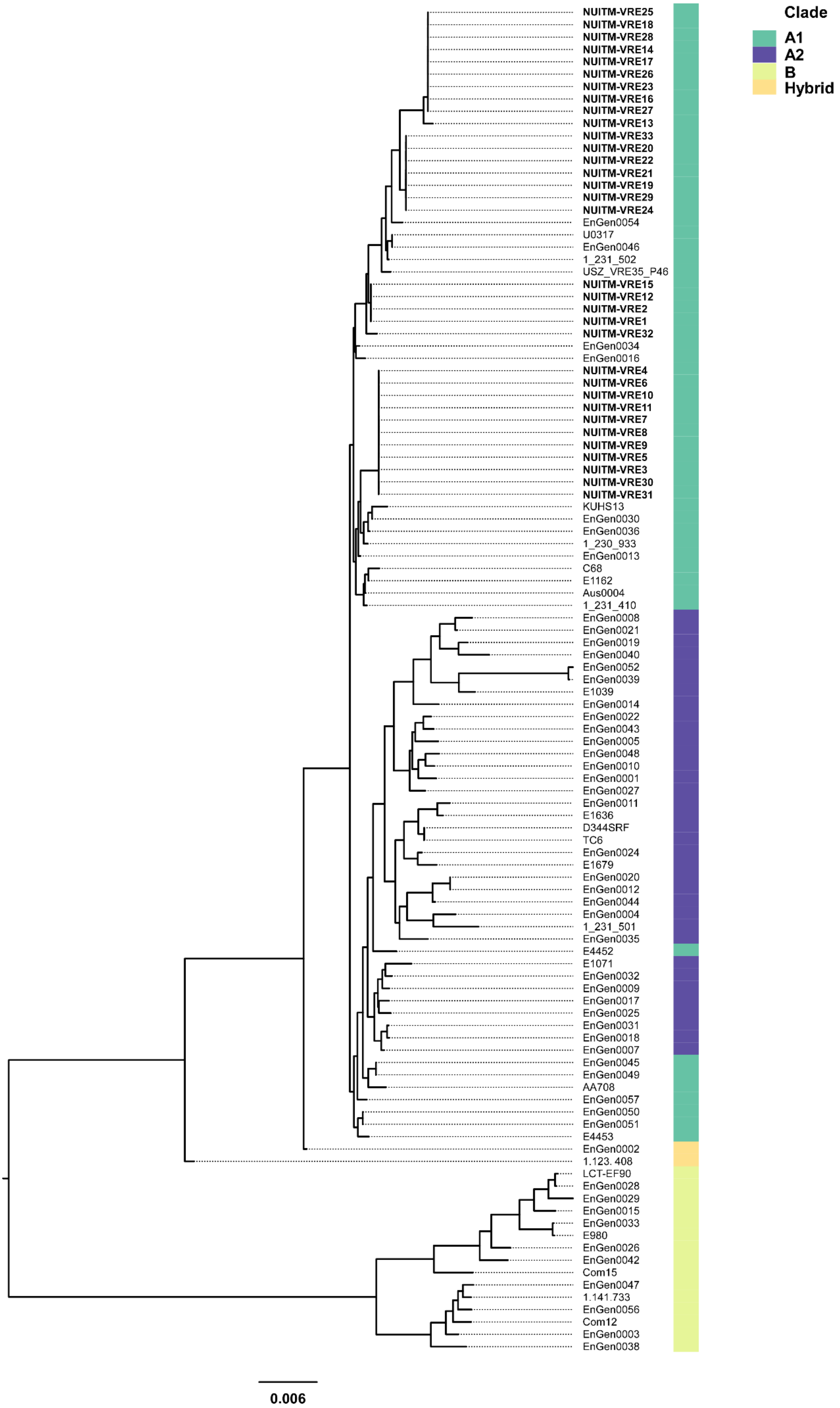
Clade
A1
A2
B
Hybrid
NUITM-VRE25
NUITM-VRE18
NUITM-VRE28
NUITM-VRE14
NUITM-VRE17
NUITM-VRE26
NUITM-VRE23
NUITM-VRE16
NUITM-VRE27
NUITM-VRE13
NUITM-VRE33
NUITM-VRE20
NUITM-VRE22
NUITM-VRE21
NUITM-VRE19
NUITM-VRE29
NUITM-VRE24
EnGen0054
U0317
EnGen0046
1_231_502
USZ_VRE35_P46
NUITM-VRE15
NUITM-VRE12
NUITM-VRE2
NUITM-VRE1
NUITM-VRE32
EnGen0034
EnGen0016
NUITM-VRE4
NUITM-VRE6
NUITM-VRE10
NUITM-VRE11
NUITM-VRE7
NUITM-VRE8
NUITM-VRE9
NUITM-VRE5
NUITM-VRE3
NUITM-VRE30
NUITM-VRE31
KUHS13
EnGen0030
EnGen0036
1_230_933
EnGen0013
C68
E1162
Aus0004
1_231_410
EnGen0008
EnGen0021
EnGen0019
EnGen0040
EnGen0052
EnGen0039
E1039
EnGen0014
EnGen0022
EnGen0043
EnGen0005
EnGen0048
EnGen0010
EnGen0001
EnGen0027
EnGen0011
E1636
D344SRF
TC6
EnGen0024
E1679
EnGen0020
EnGen0012
EnGen0044
EnGen0004
1_231_501
EnGen0035
E4452
E1071
EnGen0032
EnGen0009
EnGen0017
EnGen0025
EnGen0031
EnGen0018
EnGen0007
EnGen0045
EnGen0049
AA708
EnGen0057
EnGen0050
EnGen0051
E4453
EnGen0002
1.123. 408
LCT-EF90
EnGen0028
EnGen0029
EnGen0015
EnGen0033
E980
EnGen0026
EnGen0042
Com15
EnGen0047
1.141.733
EnGen0056
Com12
EnGen0003
EnGen0038
0.006

**Figure S2. Clade analysis of 33 VRE strains detected in Vietnamese rivers.**

Clade analysis of 33 VRE strains detected in Vietnamese rivers was performed by core genome analysis based on the report of Lebreton *et al* using Roary[11,78]. The conditions were performed with Blastp=95% and Coverage=99%. The bold text indicates the strains detected in this study, while the others are previously reported strains. Green panels indicate clade A1, purple panels indicate clade A2, yellow-green panels indicate clade B, and orange panels indicate hybrid clades.

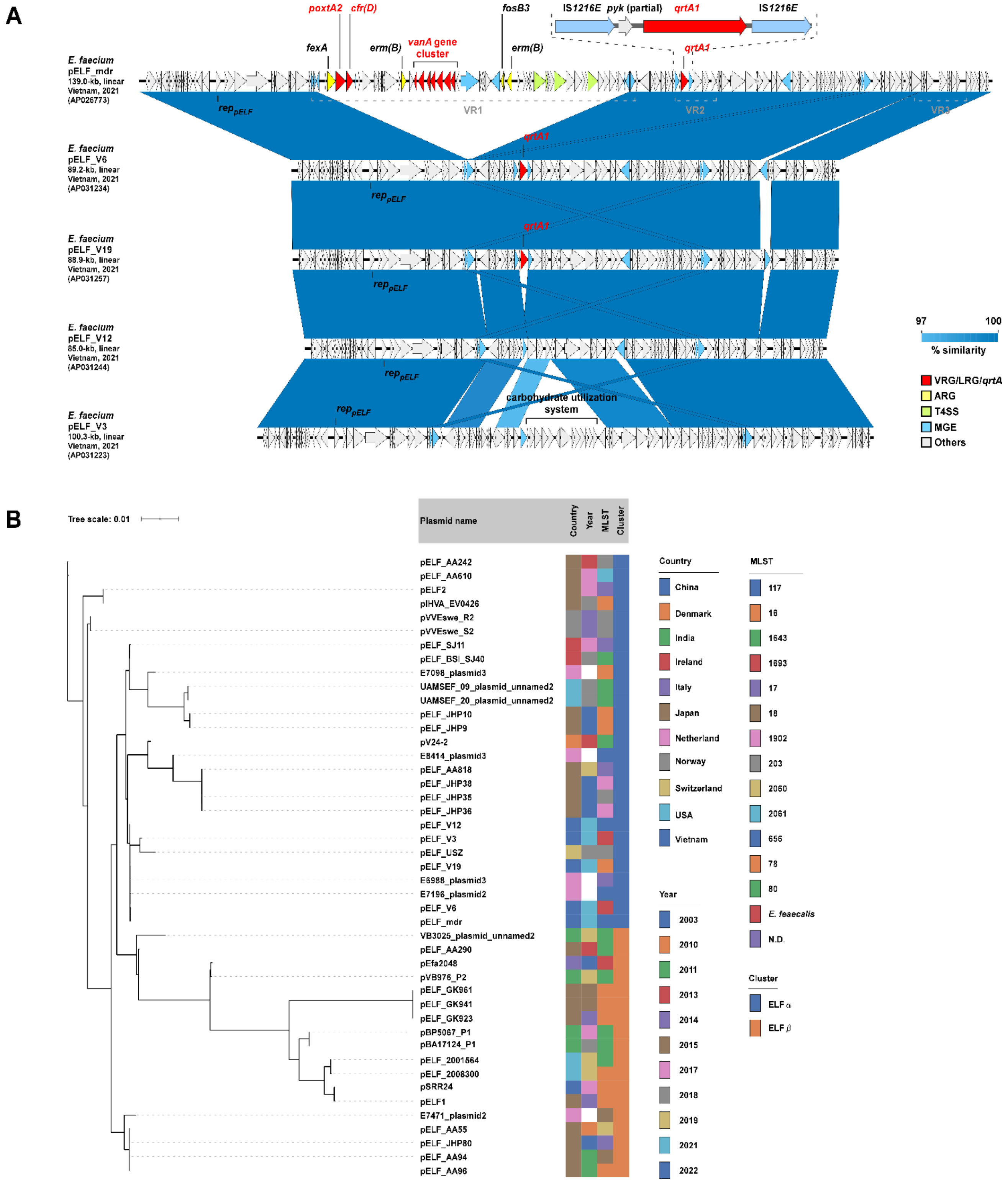


**Figure S3. Comparison of the genetic structure of pELF-type linear plasmids detected from Vietnamese rivers and core plasmid gene analysis.**

(A) A structural comparison of pELF-type linear plasmids possessed by NUITM-VRE1, VRE3, VRE6, VRE12 and VRE19 isolated from Vietnamese rivers was conducted. Red panels show vancomycin, linezolid, and *qrtA1* genes, yellow panels show other ARGs, green panels show Type 4 Secretion system, and light blue panels show MGE genes. (B) Phylogenetic analysis of 45 pELF-type linear plasmids, including pELF_mdr, was performed in Roary based on 56 plasmid core genes defined at 80% BLASTp identity and 99% alignment coverage[78], and the county and year of isolation, multilocus sequence type for *E. faecium*, and ELF_cluster are shown[30].

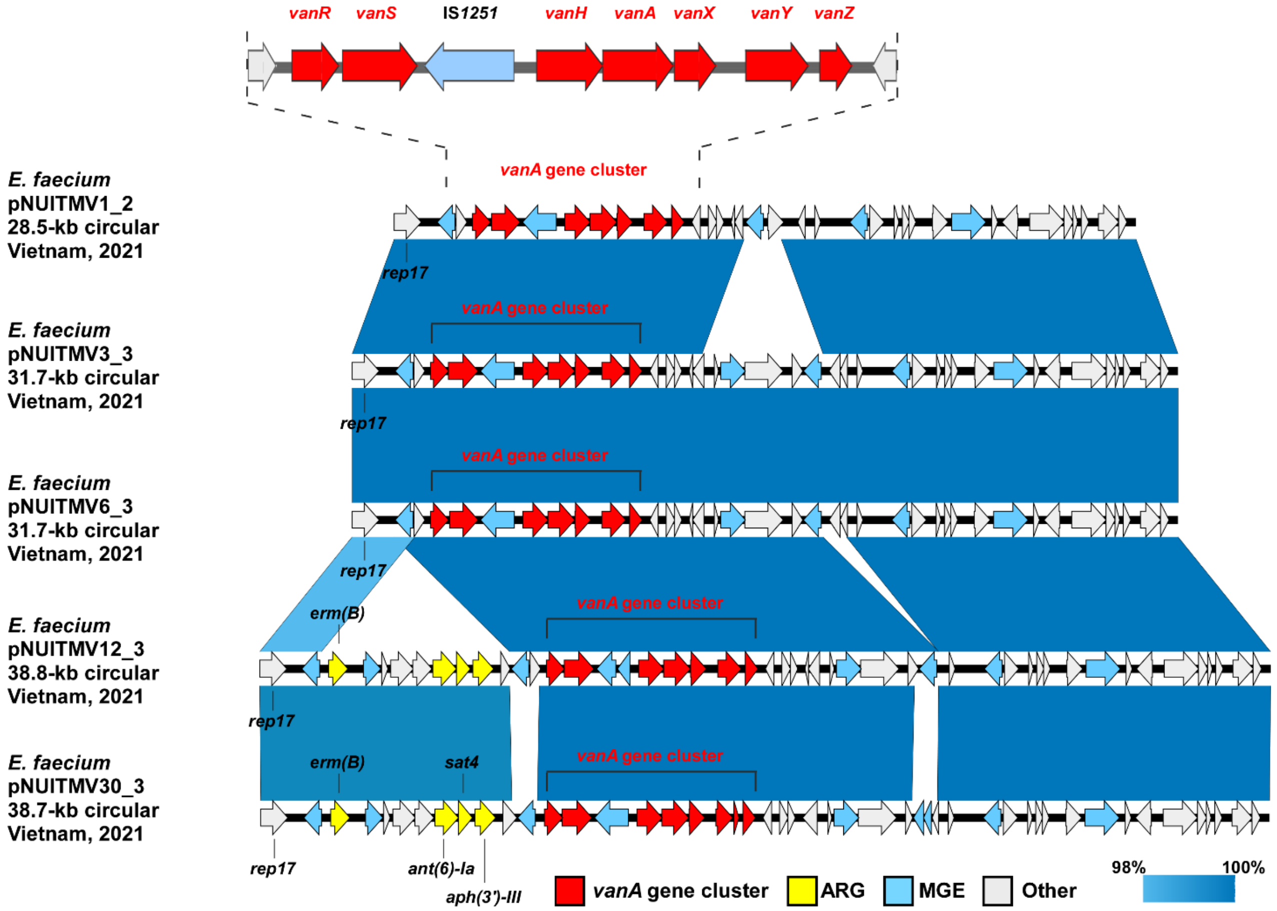


**Figure S4.** **Comparison of the genetic structure of the VanA-carrying circular plasmids harbored by NUITM-VREs.**

A structural comparison of VanA-carrying circular plasmids possessed by NUITM-VREs isolated from Vietnamese rivers was conducted. Red panels show *vanA* gene cluster, yellow panels show other ARGs, and light blue panels show MGE genes.

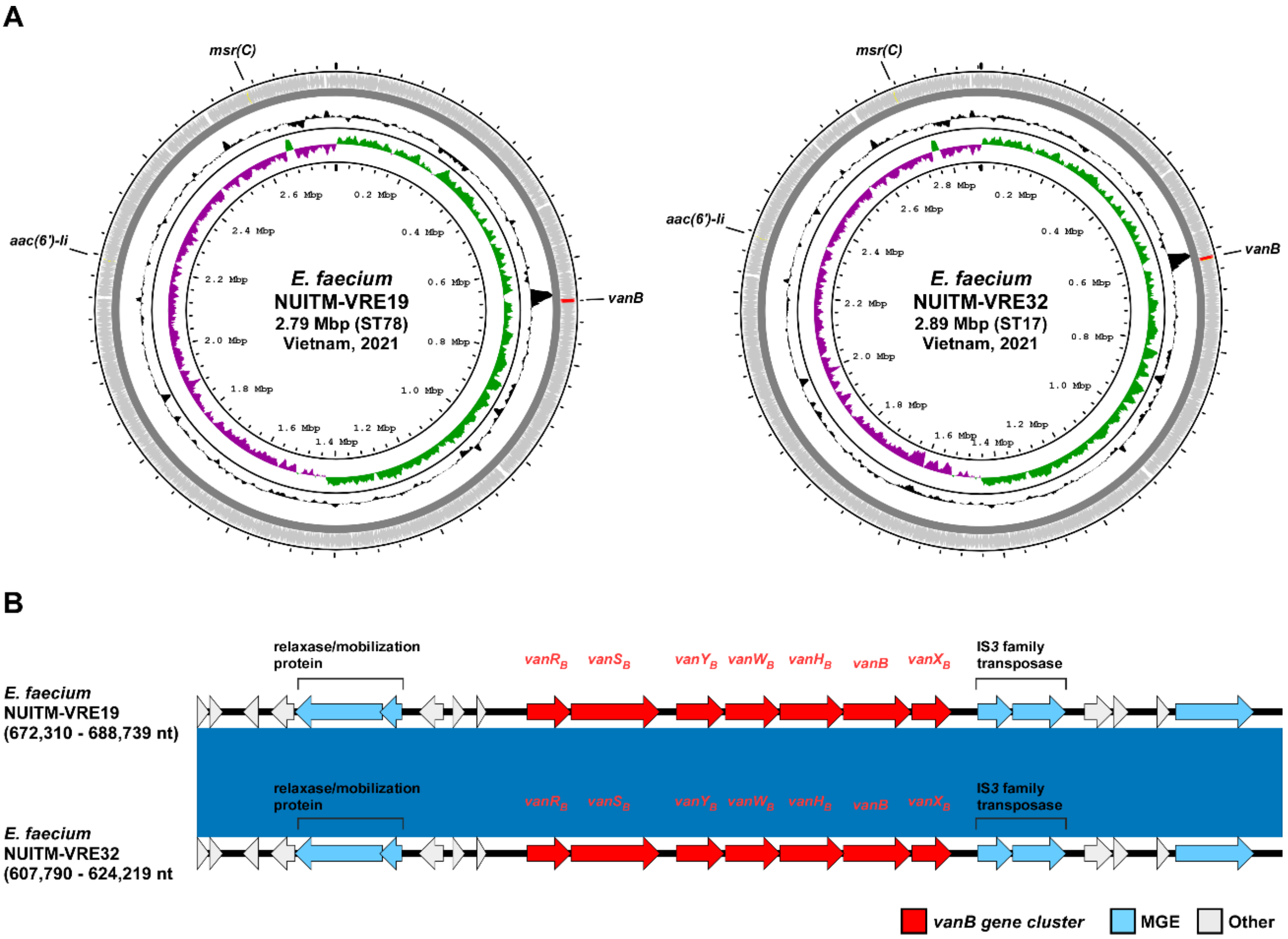


**Figure S5. Comparison of the genetic structure of the VanB-carrying Tn*1549*-like elements chromosomes of NUITM-VREs.**

Visualization of the chromosomes of VanB-carrying NUITM-VRE isolates and comparison of the genomic regions surrounding *vanB* gene cluster. Red panels show *vanB* gene cluster, and light blue panels show MGE genes (recombinases).

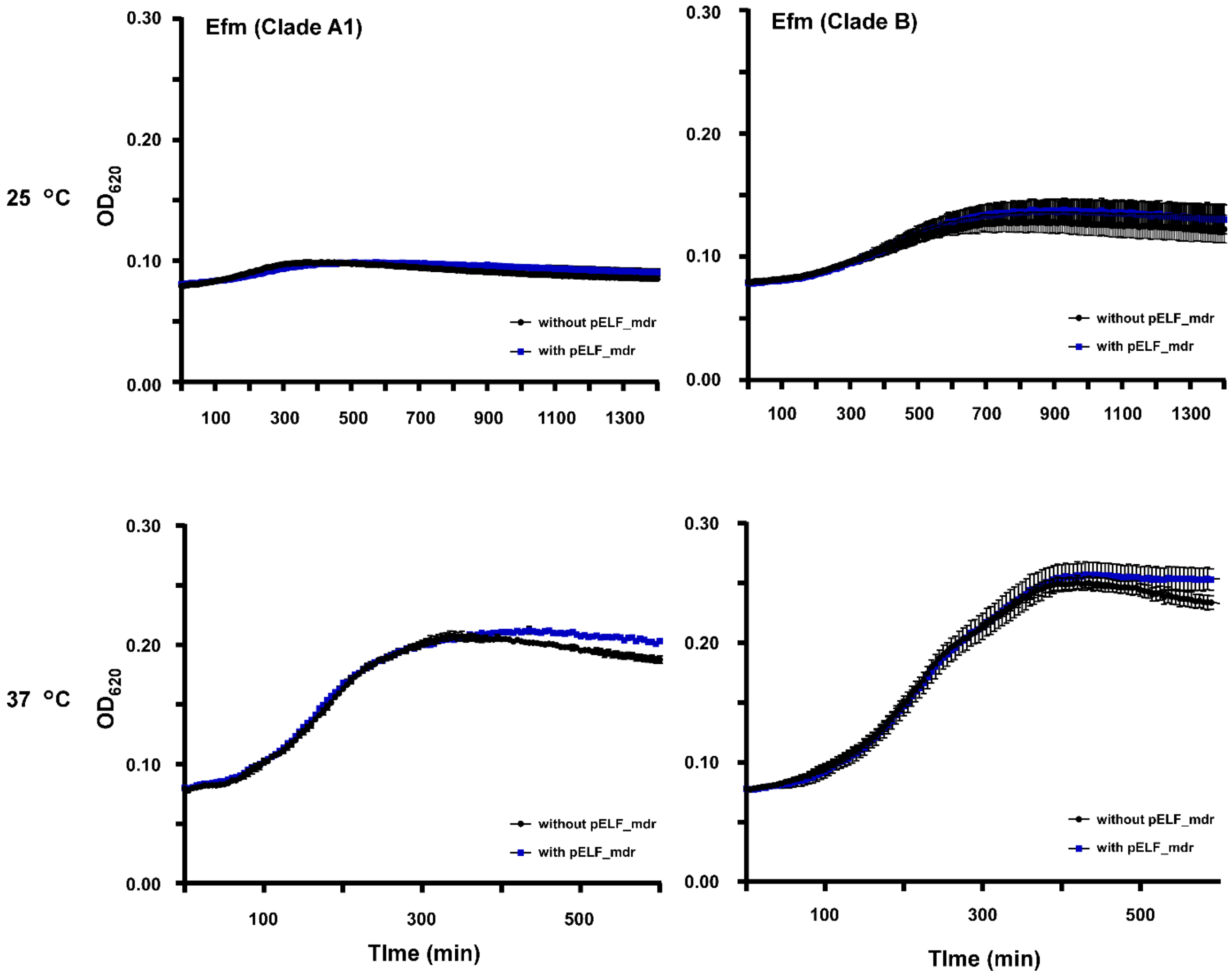


**Figure S6. Growth curve analysis of pELF_mdr-carrying *E. faecium* transconjugants on R2A medium.**

pELF_mdr-carrying clade A1 or clade B *E. faecium* transconjugants were inoculated on an R2A medium mimicking an aquatic environment, and growth curve were measured at $OD_{620}$. Blue lines indicate pELF_mdr-carrying strains, and black lines indicate non-carrying strains.

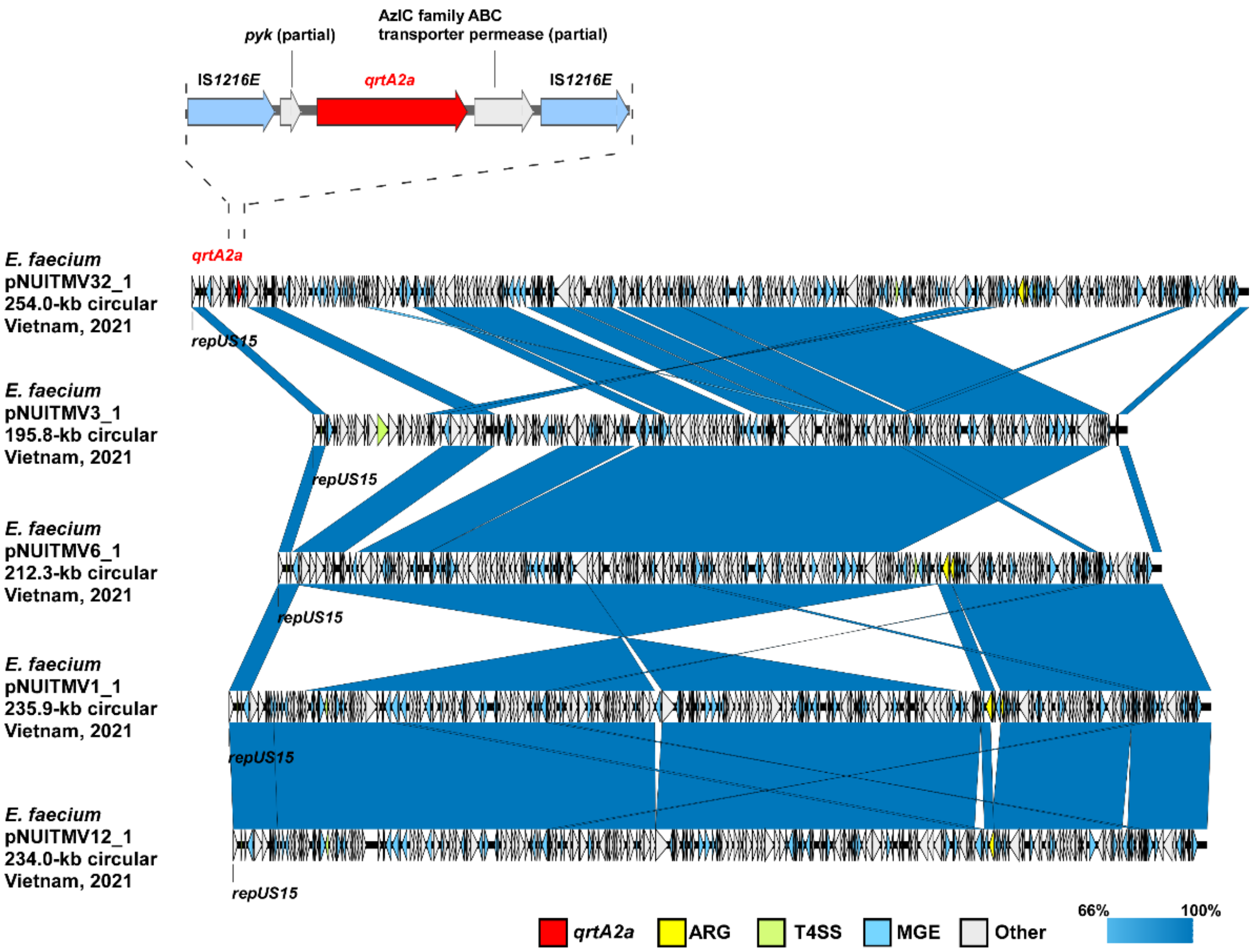


**Figure S7. Comparative analysis of megaplasmids possessed by NUITM-VREs.**

Comparative analysis of megaplasmids possessed by NUITM-VRE32, where *qrtA2a* localizes, and megaplasmids possessed by other NUITM-VREs was performed. Red arrows represent *qrtA2a* gene, yellow arrow represents ARG, green arrow represents T4SS, and light blue arrows represent MGEs.

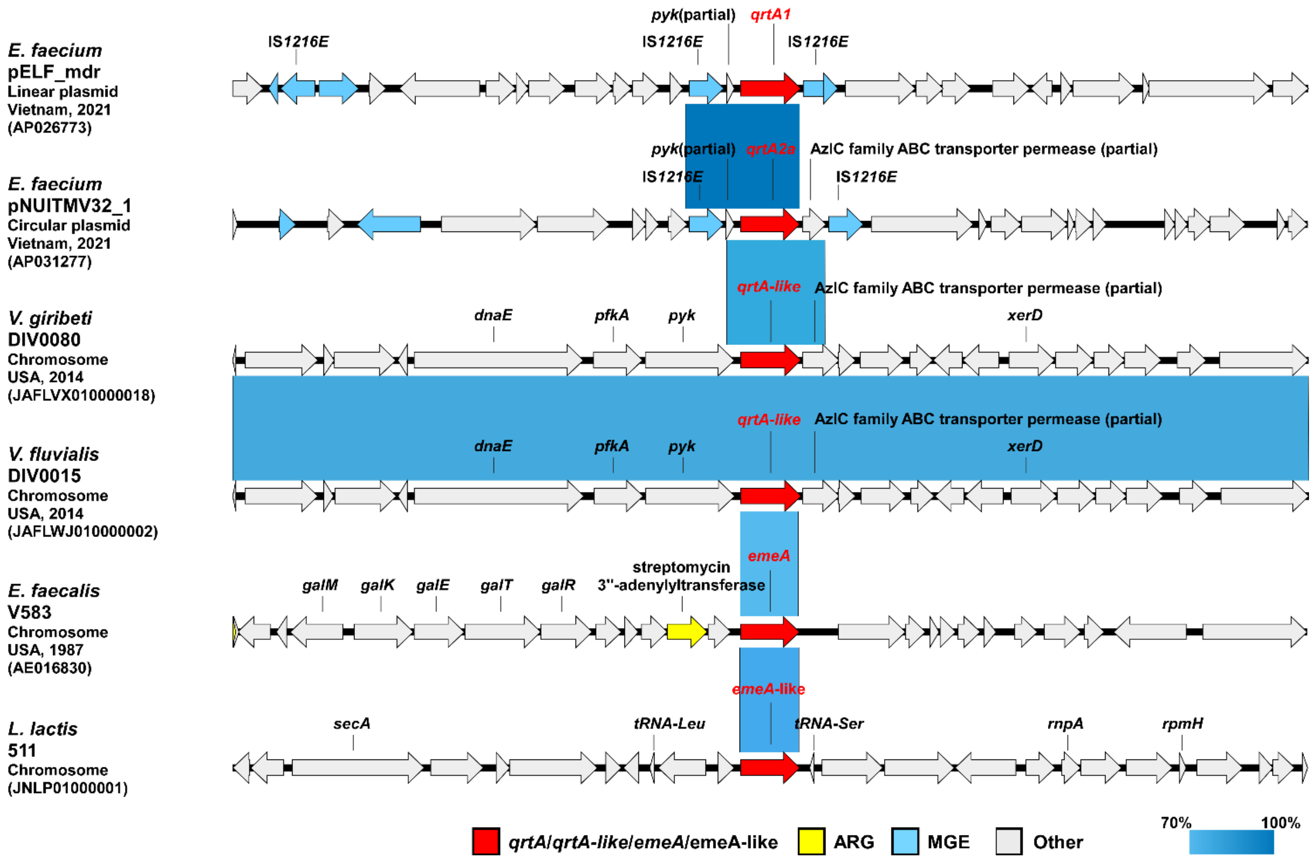


**Figure S8. Comparative analysis of the surrounding structures of MFS-type transporter genes, including *qrtA*.**

Comparative analysis of the structure around the gene of MFS-type transporters, including *qrtA*, was performed. Red arrows represent *qrtA*, *qrtA*-like from *V. giribeti* and *V. fluvialis*, *emeA*, and *emeA*-like genes, yellow arrow represents ARG, and light blue arrows represent MGEs.

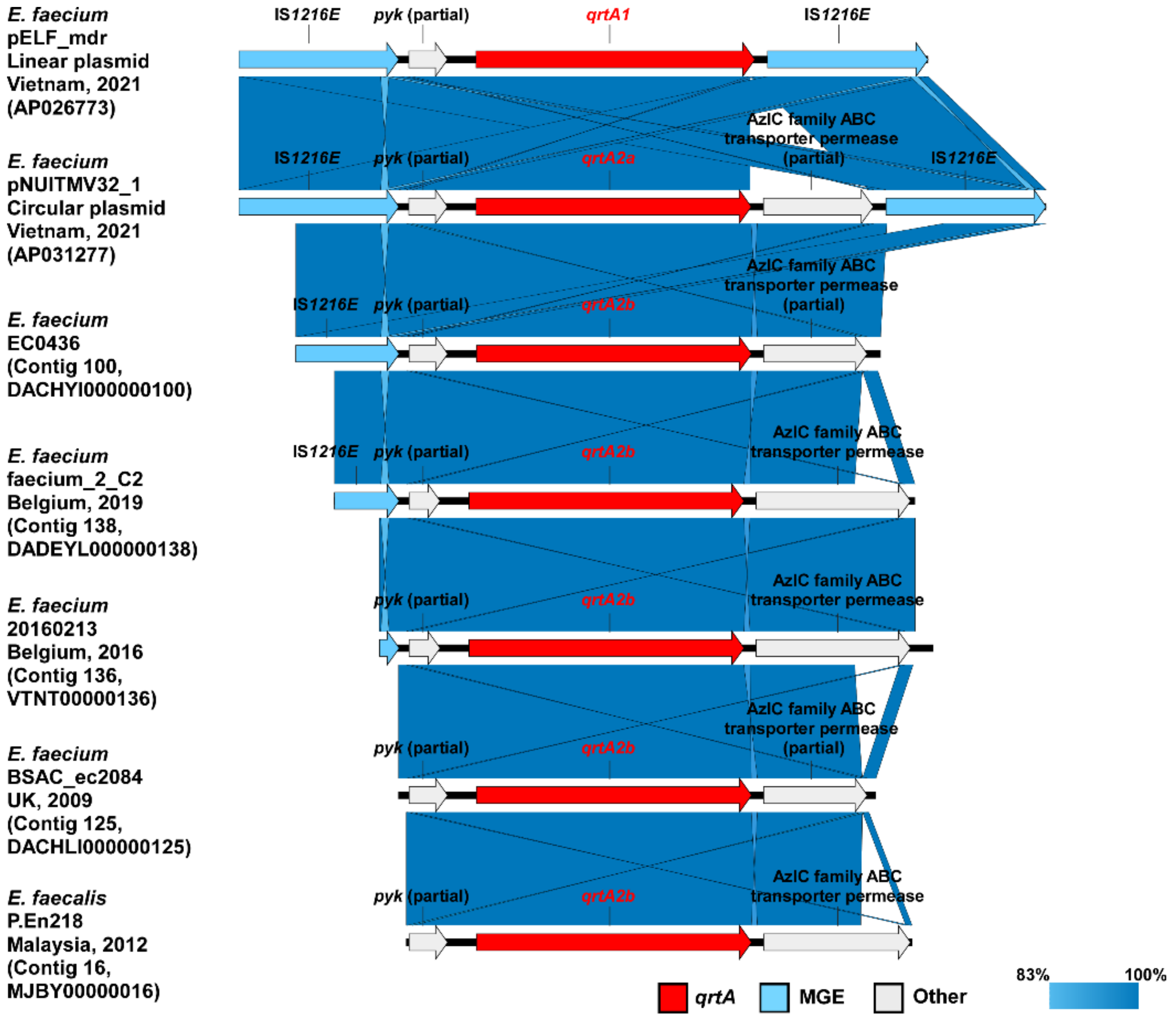


**Figure S9.** **Comparative analysis of the surrounding structure of the *qrtA* gene harbored by enterococci.**

Comparative analysis of the *qrtA* gene structure was performed for NUITM-VRE1, VRE32, and *qrtA*-positive enterococcal isolates identified by BLAST search. Red arrows indicate *qrtA* and variant genes, and light blue arrows indicate MGEs.

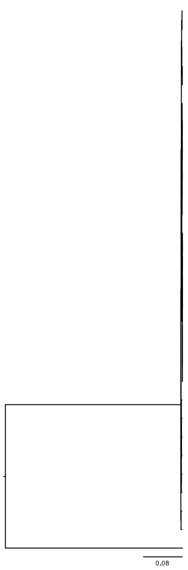


| Isolate | Species | Clade | MLST | Source | Country | Year | ARG | Plasmid | Accession no. |
|---|---|---|---|---|---|---|---|---|---|
| NUITM-VRE1 | *Enterococcus faecium* | A | ST117 | Environment | Vietnam | 2021 | *aac(6')-Ii, cat, erm(B), fexA, fosB3, msr(C), tet(L), tet(M), cfr(D), poxtA, vanA, qrtA1* | rep1, rep11a, rep17, rep18b, rep2, repUS15, repUS47, $rep_{pELF}$ | BTRX01000000 |
| NUITM-VRE2 | *Enterococcus faecium* | A | ST117 | Environment | Vietnam | 2021 | *aac(6')-Ii, cat, erm(B), fexA, fosB3, msr(C), tet(L), tet(M), cfr(D), poxtA, vanA, qrtA1* | rep1, rep11a, rep17, rep18b, rep2, repUS15, repUS47, $rep_{pELF}$ | BTSI01000000 |
| NUITM-VRE32 | *Enterococcus faecium* | A | ST17 | Environment | Vietnam | 2021 | *aac(6')-Ii, erm(B), msr(C), tet(L), vanB, qrtA2a* | rep11a, rep18b, rep2, rep22, repUS12, repUS15 | BTSL01000000 |
| 20160213 | *Enterococcus faecium* | A | ST17 | *Homo sapiens* | Belgium | 2016 | *aac(6')-aph(2''), aac(6')-Ii, aph(3')-III, erm(B), msr(C), vanB, qrtA2b* | rep11a, rep2, repUS15, repUS7, $rep_{pELF}$ | VTNT00000000 |
| faecium_2_C2 | *Enterococcus faecium* | A | ST17 | *Homo sapiens* | Belgium | 2019 | *aac(6')-aph(2''), aac(6')-Ii, aph(3')-III, erm(B), msr(C), vanB, qrtA2b* | rep11a, rep2, repUS15, repUS7, $rep_{pELF}$ | DADEYL000000000 |
| NUITM-VRE19 | *Enterococcus faecium* | A | ST78 | Environment | Vietnam | 2021 | *aac(6')-aph(2''), aac(6')-Ii, ant(6)-Ia, erm(B), msr(C), tet(L), tet(M), vanB, qrtA1* | rep14a, rep18b, rep2, repUS12, repUS15, repUS43, repUS7, $rep_{pELF}$ | BTRW01000000 |
| NUITM-VRE20 | *Enterococcus faecium* | A | ST78 | Environment | Vietnam | 2021 | *aac(6')-aph(2''), aac(6')-Ii, ant(6)-Ia, erm(B), msr(C), tet(L), tet(M), vanB, qrtA1* | rep14a, rep18b, rep2, repUS12, repUS15, repUS43, repUS7, $rep_{pELF}$ | BTRY01000000 |
| NUITM-VRE21 | *Enterococcus faecium* | A | ST78 | Environment | Vietnam | 2021 | *aac(6')-Ii, ant(6)-Ia, erm(B), msr(C), tet(L), tet(M), vanB, qrtA1* | rep14a, rep18b, rep2, repUS12, repUS43, repUS7, $rep_{pELF}$ | BTRZ01000000 |
| NUITM-VRE33 | *Enterococcus faecium* | A | ST78 | Environment | Vietnam | 2021 | *aac(6')-aph(2''), aac(6')-Ii, ant(6)-Ia, erm(B), msr(C), tet(L), tet(M), vanB, qrtA1* | rep14a, rep18b, rep2, repUS12, repUS15, repUS43, repUS7, $rep_{pELF}$ | BTSM01000000 |
| NUITM-VRE24 | *Enterococcus faecium* | A | ST78 | Environment | Vietnam | 2021 | *aac(6')-aph(2''), aac(6')-Ii, ant(6)-Ia, erm(B), msr(C), tet(L), tet(M), vanB, qrtA1* | rep14a, rep18b, rep2, repUS12, repUS15, repUS43, repUS7, $rep_{pELF}$ | BTSC01000000 |
| NUITM-VRE29 | *Enterococcus faecium* | A | ST78 | Environment | Vietnam | 2021 | *aac(6')-aph(2''), aac(6')-Ii, ant(6)-Ia, erm(B), msr(C), tet(L), tet(M), vanB, qrtA1* | rep14a, rep18b, rep2, repUS12, repUS15, repUS43, repUS7, $rep_{pELF}$ | BTSH01000000 |
| NUITM-VRE22 | *Enterococcus faecium* | A | ST78 | Environment | Vietnam | 2021 | *aac(6')-aph(2''), aac(6')-Ii, ant(6)-Ia, erm(B), msr(C), tet(L), tet(M), vanB, qrtA1* | rep14a, rep18b, rep2, repUS12, repUS15, repUS43, repUS7, $rep_{pELF}$ | BTSA01000000 |
| NUITM-VRE14 | *Enterococcus faecium* | A | ST80 | Environment | Vietnam | 2021 | *aac(6')-Ii, dfrG, msr(C), vanA, qrtA1* | rep14a, rep17, rep29, repUS15, $rep_{pELF}$ | BTRR01000000 |
| NUITM-VRE17 | *Enterococcus faecium* | A | ST80 | Environment | Vietnam | 2021 | *aac(6')-Ii, dfrG, msr(C), vanA, qrtA1* | rep14a, rep29, repUS15, $rep_{pELF}$ | BTRU01000000 |
| NUITM-VRE26 | *Enterococcus faecium* | A | ST80 | Environment | Vietnam | 2021 | *aac(6')-Ii, dfrG, msr(C), vanA, qrtA1* | rep14a, repUS15, $rep_{pELF}$ | BTSE01000000 |
| NUITM-VRE16 | *Enterococcus faecium* | A | ST80 | Environment | Vietnam | 2021 | *aac(6')-Ii, dfrG, msr(C), vanA, qrtA1* | rep14a, rep17, rep29, repUS15, $rep_{pELF}$ | BTRT01000000 |
| NUITM-VRE23 | *Enterococcus faecium* | A | ST80 | Environment | Vietnam | 2021 | *aac(6')-Ii, dfrG, msr(C), vanA, qrtA1* | rep14a, rep17, rep29, repUS15, $rep_{pELF}$ | BTSB01000000 |
| NUITM-VRE18 | *Enterococcus faecium* | A | ST80 | Environment | Vietnam | 2021 | *aac(6')-Ii, dfrG, msr(C), vanA, qrtA1* | rep14a, rep17, rep29, repUS15, $rep_{pELF}$ | BTRV01000000 |
| NUITM-VRE25 | *Enterococcus faecium* | A | ST80 | Environment | Vietnam | 2021 | *aac(6')-Ii, dfrG, msr(C), vanA, qrtA1* | rep14a, rep17, rep29, repUS15, $rep_{pELF}$ | BTSD01000000 |
| NUITM-VRE28 | *Enterococcus faecium* | A | ST80 | Environment | Vietnam | 2021 | *aac(6')-Ii, dfrG, msr(C), vanA, qrtA1* | rep14a, rep17, rep29, repUS15, $rep_{pELF}$ | BTSG01000000 |
| NUITM-VRE27 | *Enterococcus faecium* | A | ST80 | Environment | Vietnam | 2021 | *aac(6')-Ii, dfrG, msr(C), vanA, qrtA1* | rep14a, rep29, repUS15, $rep_{pELF}$ | BTSF01000000 |
| NUITM-VRE6 | *Enterococcus faecium* | A | ST1693 | Environment | Vietnam | 2021 | *aac(6')-aph(2''), aac(6')-Ii, aph(3')-III, dfrG, erm(B), lnu(B), lsa(E), msr(C), tet(M), vanA, qrtA1* | rep11a, rep14a, rep17, rep18b, rep2, repUS15, repUS43, $rep_{pELF}$ | BTST01000000 |
| NUITM-VRE11 | *Enterococcus faecium* | A | ST1693 | Environment | Vietnam | 2021 | *aac(6')-aph(2''), aac(6')-Ii, aph(3')-III, dfrG, erm(B), lnu(B), lsa(E), msr(C), tet(M), vanA, qrtA1* | rep11a, rep14a, rep17, rep18b, rep2, repUS15, repUS43, $rep_{pELF}$ | BTRO01000000 |
| NUITM-VRE7 | *Enterococcus faecium* | A | ST1693 | Environment | Vietnam | 2021 | *aac(6')-aph(2''), aac(6')-Ii, aph(3')-III, dfrG, erm(B), lnu(B), lsa(E), msr(C), tet(M), vanA, qrtA1* | rep11a, rep14a, rep17, rep18b, rep2, repUS15, repUS43, $rep_{pELF}$ | BTSU01000000 |
| NUITM-VRE9 | *Enterococcus faecium* | A | ST1693 | Environment | Vietnam | 2021 | *aac(6')-aph(2''), aac(6')-Ii, aph(3')-III, dfrG, erm(B), lnu(B), lsa(E), msr(C), tet(M), vanA, qrtA1* | rep11a, rep14a, rep17, rep18b, rep2, repUS15, repUS43, $rep_{pELF}$ | BTSW01000000 |
| NUITM-VRE10 | *Enterococcus faecium* | A | ST1693 | Environment | Vietnam | 2021 | *aac(6')-aph(2''), aac(6')-Ii, aph(3')-III, dfrG, erm(B), lnu(B), lsa(E), msr(C), tet(M), vanA, qrtA1* | rep11a, rep14a, rep17, rep18b, rep2, repUS15, repUS43, $rep_{pELF}$ | BTRN01000000 |
| NUITM-VRE8 | *Enterococcus faecium* | A | ST1693 | Environment | Vietnam | 2021 | *aac(6')-aph(2''), aac(6')-Ii, aph(3')-III, dfrG, erm(B), lnu(B), lsa(E), msr(C), tet(M), vanA, qrtA1* | rep11a, rep14a, rep17, rep18b, rep2, repUS15, repUS43, $rep_{pELF}$ | BTSV01000000 |
| BSAC_ec2084 | *Enterococcus faecium* | A | ST280 | *Homo sapiens* | UK | 2009 | *aac(6')-aph(2''), aac(6')-Ii, ant(9)-Ia, erm(A), erm(T), msr(C), qrtA2b* | rep14a, rep14b, rep18b, rep2, rep29, repUS15, repUS43 | DACHLI000000000 |
| EC0436 | *Enterococcus faecium* | A | ST262 | N/A | N/A | N/A | *aac(6')-aph(2''), aac(6')-Ii, ant(6)-Ia, ant(9)-Ia, dfrG, erm(A), erm(B), lnu(B), lsa(E), msr(C), tet(L), tet(M), qrtA2b* | rep14a, rep14b, rep2, repUS15, repUS43 | DACHYI000000000 |
| P.En218 | *Enterococcus faecalis* | N/A | ST765 | *Homo sapiens* | Malaysia | 2012 | *aac(6')-aph(2''), ant(6)-Ia, ant(9)-Ia, aph(3')-III, cat, dfrG, erm(A), erm(B), fexA, lnu(B), lsa(A), lsa(E), str, tet(L), tet(M), optrA, qrtA2b* | rep7a, rep9a, repUS11, repUS43 | MJBY00000000 |

**Figure S10 Phylogenetic analysis of enterococci harboring the *qrtA* gene.**

Phylogenetic tree of *qrtA*-positive enterococcal isolates; bacterial species, clade, multilocus sequence type (ST), source, country and year of isolation, the presence of *qrtA* alleles (*qrtA1*, *qrtA2* and *qrtA*-like, highlighted in blue), co-located ARGs, pELF-type linear plasmids rep ($rep_{pELF}$) and other plasmid *rep* genes, and accession numbers are shown.

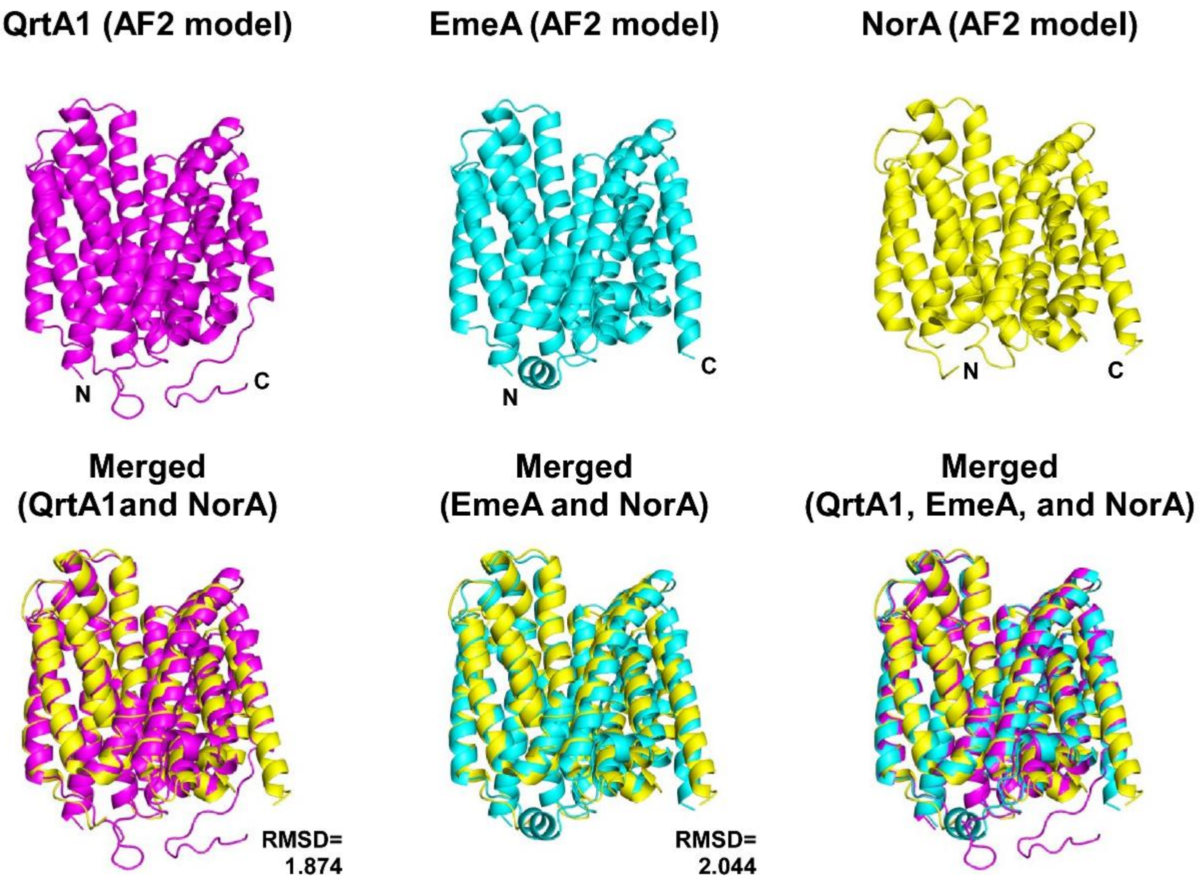


**Figure S11. Structural analysis of QrtA1, EmeA, and NorA based on AlfaFold2.**

Protein structure prediction using amino acid sequences using AlphaFold v2.3.1 was performed on the Galaxy Europe server with default parameters. AlphaFold-predicted and experimental structures were visualized and compared using the PyMOL software v2.5.8.

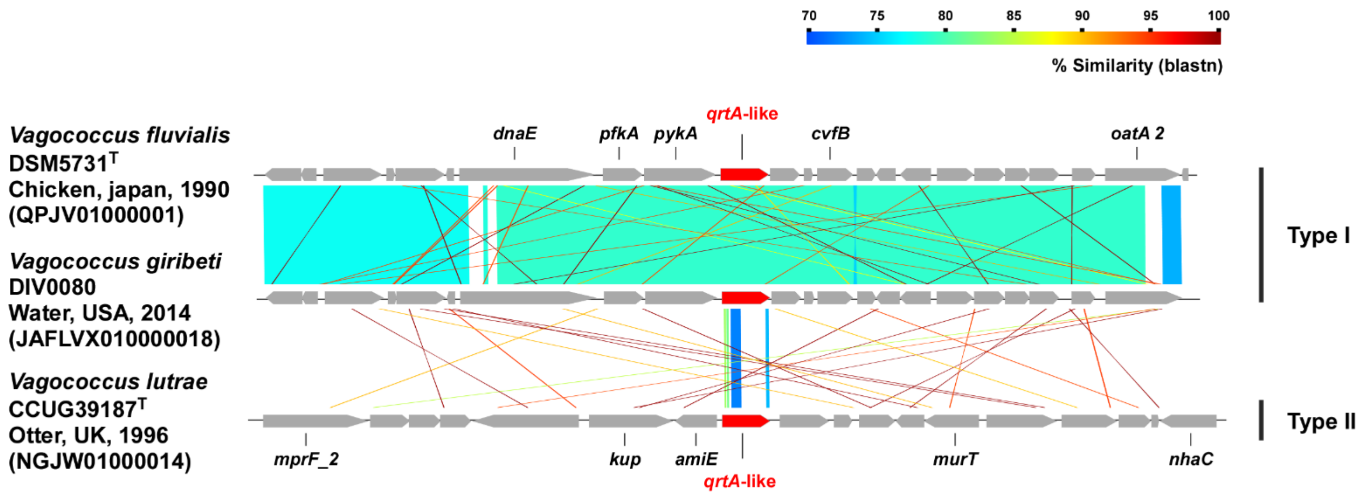


**Figure 12. Comparison of the surrounding genetic structures of the *qrtA*-like gene.**

Comparison of the surrounding genetic structures of the qrtA-like gene on the chromosomes of Vogococcus species was performed and illustrated. The bacterial genomes used in this study are as follows: *V. fluvialis* DSM 5731r (Accession No. QPJV01000001), *V. giribeti* DIV0080 (JAFLVX010000018), and *V. lutrae* CCUG 39187r (NGJW01000014).

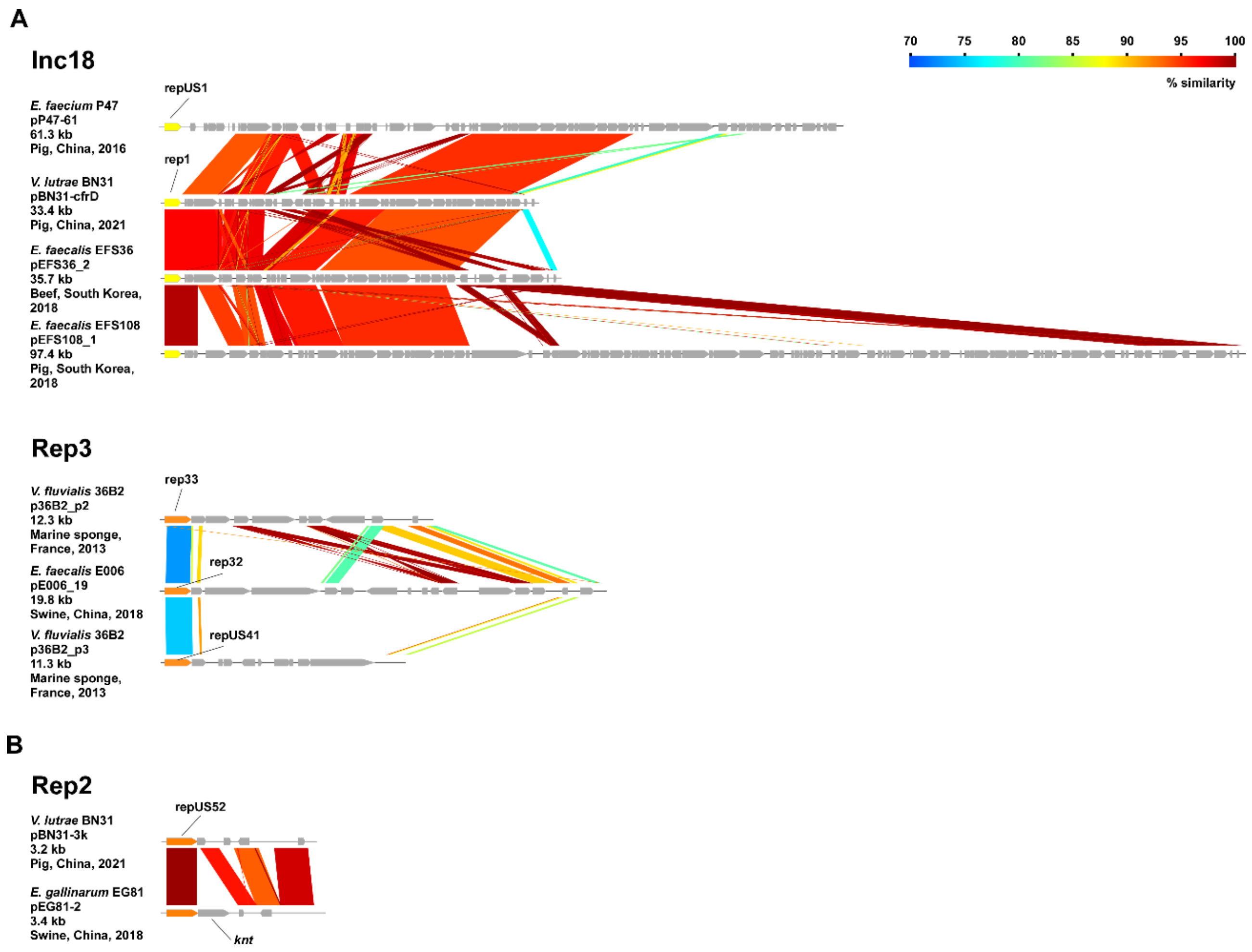


**Figure S13. Comparative analysis of circular plasmids held by vagococci and enterococci.**

(A) Comparison of vagococci and enterococci plasmids harboring Inc18-type and Rep3-type reps was performed and illustrated. The plasmids used are as follows; *V. lutrae* BN31 pBN31-cfrD (CP081834); *E. faecium* P47 pP47-61 (CP091102); *E. faecalis* EFS36 pEFS36_2 (CP085293); *E. faecalis* EFS108 pEFS108_1 (CP085295); *V. fluvialis* 36B2 p36B2_p2 (CP081463); *E. faecalis* E006 pE006_19 (CP082233); *V. fluvialis* 36B2 p36B2_p3 (CP081464). (B) Comparative analysis of plasmids of *E. gallinarum*, a non-*E. faecium*/*E. faecalis*, and *V. lutrae* was performed and illustrated. The plasmids used are as follows; *V. lutrae* BN31 pBN31-3k (NZ_CP081836); *E. gallinarum* EG81 pEG81-2 (NZ_CP050818).

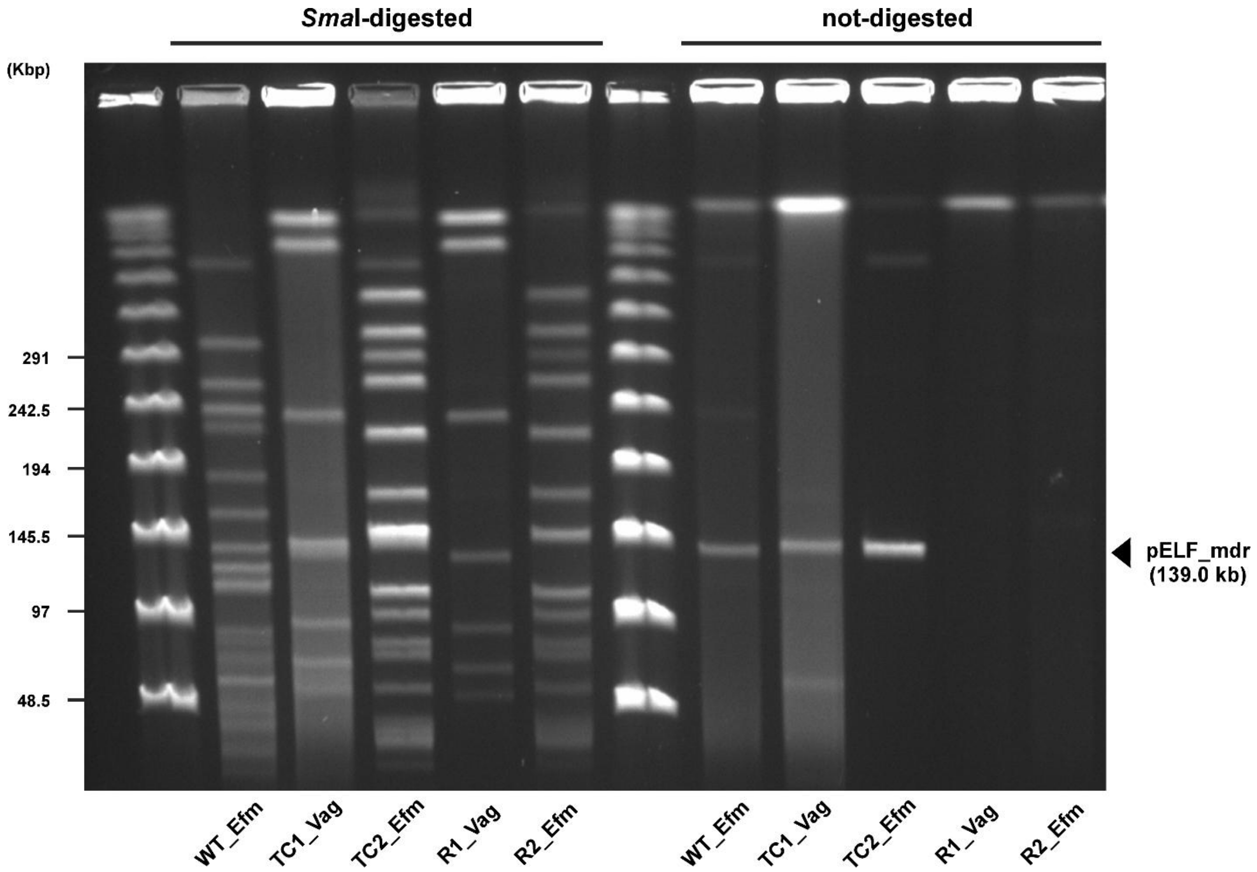


**Figure S14. PFGE analysis of pELF_mdr-carrying transconjugants between *Vagococcus* and *Enterococcus.***

Filter mating conjugation experiments were performed with *E. faecium* NUITM-VRE1 (wild-type) as a donor strain, *V. giribeti* DIV0080RF (*V. giribeti*) as the first recipient strain, and *E. faecium* BM4105SS as the second recipient strain; transconjugants were subjected to pulsed-fired gel electrophoresis. Total DNA treated with the restriction enzyme (*Sma*I-digested) was used to confirm the chromosomal type, and total DNA without S1-nuclease treatment (not-digested) was used to confirm pELF_mdr linear plasmids. The bands of pELF_mdr are noted with an arrowhead. WT_Efm: NUITM-VRE1; TC1_Vag: DIV0080RF/pELF_mdr; TC2_Efm: BM4105SS/pELF_mdr; R1_Vag: DIV0080RF; R2_Efm: BM4105SS.

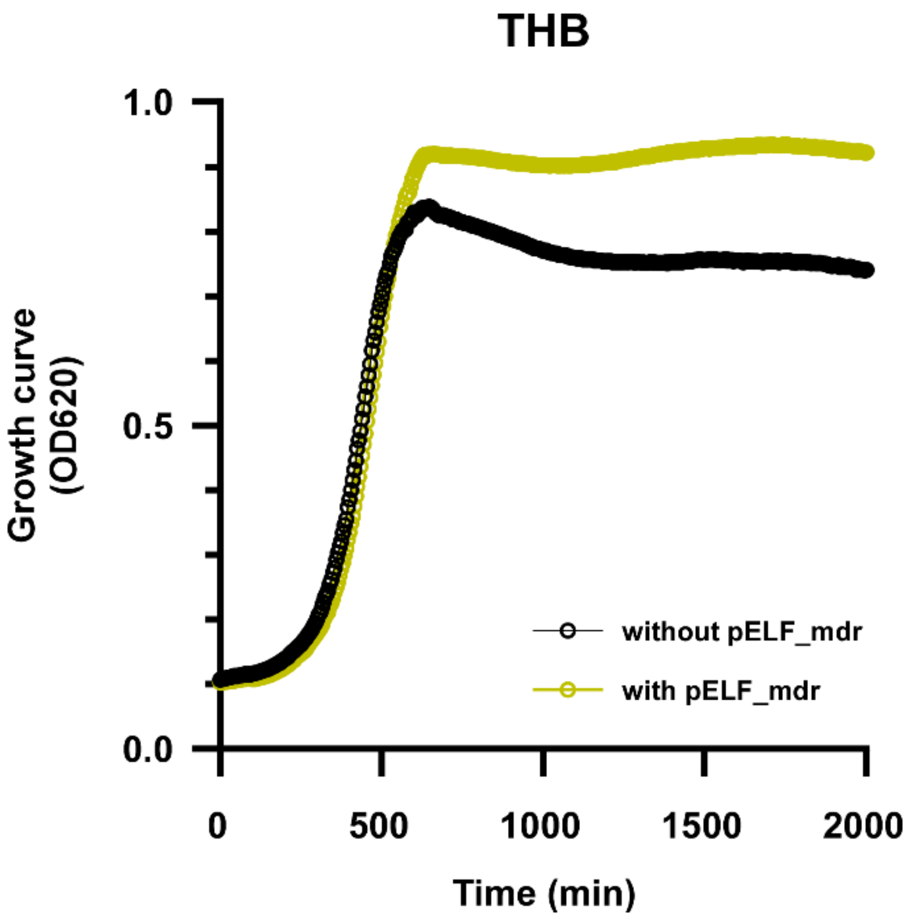


**Figure S15. Growth curve analysis of pELF_mdr-carrying transconjugant of *Vagococcus giribeti*.**

Growth curves of *Vagococcus giribeti* (DIV0080RF) and its pELF/mdr transconjugant (DIV0080RF/pELF_mdr) were measured at $OD_{620}$ in THB medium at 37°C. Black circles and lines indicate wild type (DIV0080RF), yellow circles and lines indicate pELF_mdr transconjugant (DIV0080RF/pELF_mdr).
